# Keratin 19 interacts with GSK3β to regulate its nuclear accumulation and degradation of cyclin D3

**DOI:** 10.1101/2021.07.28.454205

**Authors:** Pooja Sharma, Sarah Tiufekchiev, Victoria Lising, Seung Woo Chung, Jung Soo Suk, Byung Min Chung

## Abstract

Cyclin D3 regulates the G1/S transition and is frequently overexpressed in several cancer types including breast cancer, where it promotes tumor progression. Here, we show that a cytoskeletal protein keratin 19 (K19) physically interacts with a serine/threonine kinase GSK3β and prevents GSK3β-dependent degradation of cyclin D3. The absence of K19 allowed active GSK3β to accumulate in the nucleus and degrade cyclin D3. Specifically, the head domain of K19 was required to sustain inhibitory phosphorylation of GSK3β Ser9, prevent nuclear accumulation of GSK3β, and maintain cyclin D3 levels and cell proliferation. K19 was found to interact with GSK3β and K19-GSK3β interaction was mapped out to require Ser10 and Ser35 residues on the head domain of K19. Unlike wildtype K19, S10A and S35A mutants failed to maintain total and nuclear cyclin D3 levels and induce cell proliferation. Finally, we show that the K19-GSK3β-cyclin D3 pathway affected sensitivity of cells towards inhibitors to cyclin dependent kinase 4 and 6 (CDK4/6). Overall, these findings establish a role for K19 in the regulation of GSK3β-cyclin D3 pathway and demonstrate a potential strategy for overcoming resistance to CDK4/6 inhibitors.

## Introduction

Cyclins play a major role in the cell cycle progression. During the G1 phase, D-type cyclins dimerize with and activate cyclin dependent kinase 4 or 6 (CDK4/6) which then phosphorylates tumor suppressor retinoblastoma protein (Rb) (Musgrove *et al.*, 2011). Prior to phosphorylation, Rb is bound to a transcription factor E2F1 and prevents activity of E2F1 for transition from G1 to S phase (Giacinti and Giordano, 2006). Hyperphosphorylation of Rb relieves inhibition of E2F1 activity and hence induces progression to the S phase in the cell cycle (Bartkova *et al.*, 1998).

Cyclin D3 is one of three cyclin Ds, and it is overexpressed in several human cancers including breast cancer (Filipits *et al.*, 2002; Sicinska *et al.*, 2003; Weroha *et al.*, 2006; Zhang *et al.*, 2011; Chi *et al.*, 2015; Wang *et al.*, 2019). Inhibition of cyclin D3 expression in mammary tumor cells reduced cancer cell proliferation in vitro and decreased the tumor burden in vivo (Zhang *et al.*, 2011), demonstrating the oncogenic role of cyclin D3 in cancer. Inside the cell, regulation of cyclin D3 levels has been shown to involve proteasomal degradation mediated by phosphorylation of cyclin D3 at Thr283 by glycogen synthase kinase-3β (GSK3β) in acute lymphoblastic leukemia Reh cells and p38^SAPK2^ in Jurkat T cells (Casanovas *et al.*, 2004; Naderi *et al.*, 2004). However, the mechanism of cyclin D3 degradation in epithelial cancer remains unclear.

GSK3β is a multifunctional serine/threonine kinase which interacts with and phosphorylates target proteins to mediate multiple cellular processes, including proliferation, differentiation, motility and survival, apoptosis, micropinocytosis, and stress-induced autophagy (Azoulay-Alfaguter *et al.*, 2015; Albrecht *et al.*, 2020; Alao *et al.*, 2006). At the molecular level, GSK3β activity is inactivated by phosphorylation at Ser9 by kinases including Akt and Rsk (Jope and Johnson, 2004; Beurel *et al.*, 2015). Inappropriate regulation of GSK3β activity and subcellular localization contributes to the pathogenesis and progression of various diseases including non-insulin-dependent diabetes mellitus, cardiovascular disease, some neurodegenerative diseases, bipolar disorder, and cancer (Beurel *et al.*, 2015; Manning and Toker, 2017).

Here, we report a novel regulation of GSK3β activity by keratin 19 (K19). K19 belongs to a keratin family of intermediate filament proteins. Keratins are critical in maintaining structural integrity of epithelial cells and tissues, but they are also involved in other cellular processes such as proliferation and migration, especially in disease settings (Alix-Panabieres *et al.*, 2009; Sharma *et al.*, 2019b). While its normal function has not been studied in detail, K19 is expressed in the developing embryo, mature striated muscles, epithelia and epithelial stem cells (Moll *et al.*, 1982; Stone *et al.*, 2007; Petersen and Polyak, 2010). Its expression is also observed in pathological conditions including several cancer types where K19 is used as a diagnostic and prognostic marker (Alix-Panabieres *et al.*, 2009; Kabir *et al.*, 2014). Studies using hepatocellular carcinoma (Takano *et al.*, 2016), oral squamous cell carcinoma (Crowe *et al.*, 1999), breast cancer (Sharma *et al.*, 2019a), and lung cancer (Ohtsuka *et al.*, 2016) cell lines have shown that K19 promotes cancer cell proliferation. We previously used *KRT19* knockout (KO) of MCF7 breast cancer cells to identify that K19 is required for proper cell cycle progression and maintenance of levels of D-type cyclins (cyclin D1 and cyclin D3) (Sharma *et al.*, 2019a). In particular, K19 was shown to delay degradation of cyclin D3, suggesting that K19 promotes cell proliferation by stabilizing cyclin D3. However, how a cytoskeletal protein can maintain levels of cyclin D3 remained unknown.

In this study, we identify GSK3β as a keratin-interacting protein and find that K19 suppresses GSK3β activity and hinders its nuclear accumulation. GSK3β-binding by K19 required Ser10 and Ser35 residues, and K19 S10A and S35A mutants failed to maintain cyclin D3 levels. Our results reveal a novel regulatory role on GSK3β localization and activity by K19 and provide a mechanism of how a cytoskeletal protein and a signaling molecule coordinate cell proliferation. Clinically, K19-dependent regulation of GSK3β activity may be used to reduce resistance to CDK4/6 inhibitors as *KRT19* KO cells became sensitized to CDK4/6 inhibitors upon coculturing with a GSK3β inhibitor. Given that K19 expression levels are frequently elevated in various cancers, it may be used to predict the efficacy of CDK4/6 inhibitors and resistant patients may be cotreated with a GSK3β inhibitor.

## Results

### Cyclin D3 levels are downregulated by GSK3β in KRT19 KO cells

Since *KRT19* KO cells exhibit decreased cell proliferation and express decreased levels of cyclin D3 (Sharma *et al.*, 2019a), colony formation assay was performed to assess the role of cyclin D3 in cell proliferation by K19. Unlike MCF7 parental cells where overexpressing cyclin D3 did not increase colony area, overexpression of cyclin D3 significantly increased colony area compared to vector transfection in *KRT19* KO cells (Figure 1A, S1). To confirm the requirement of K19 in protein stability of cyclin D3, cyclin D3 levels were examined upon inhibition of protein synthesis using cycloheximide. Cyclin D3 levels exhibited a greater decrease in cycloheximide-treated *KRT19* KO cells compared to that of the parental control (Figure S2) (Sharma *et al.*, 2019a). Indeed, inhibiting proteasome function with MG132 markedly enhanced cyclin D3 protein levels in both of *KRT19* KO clones tested (Figure 1B).

**Figure 1.**
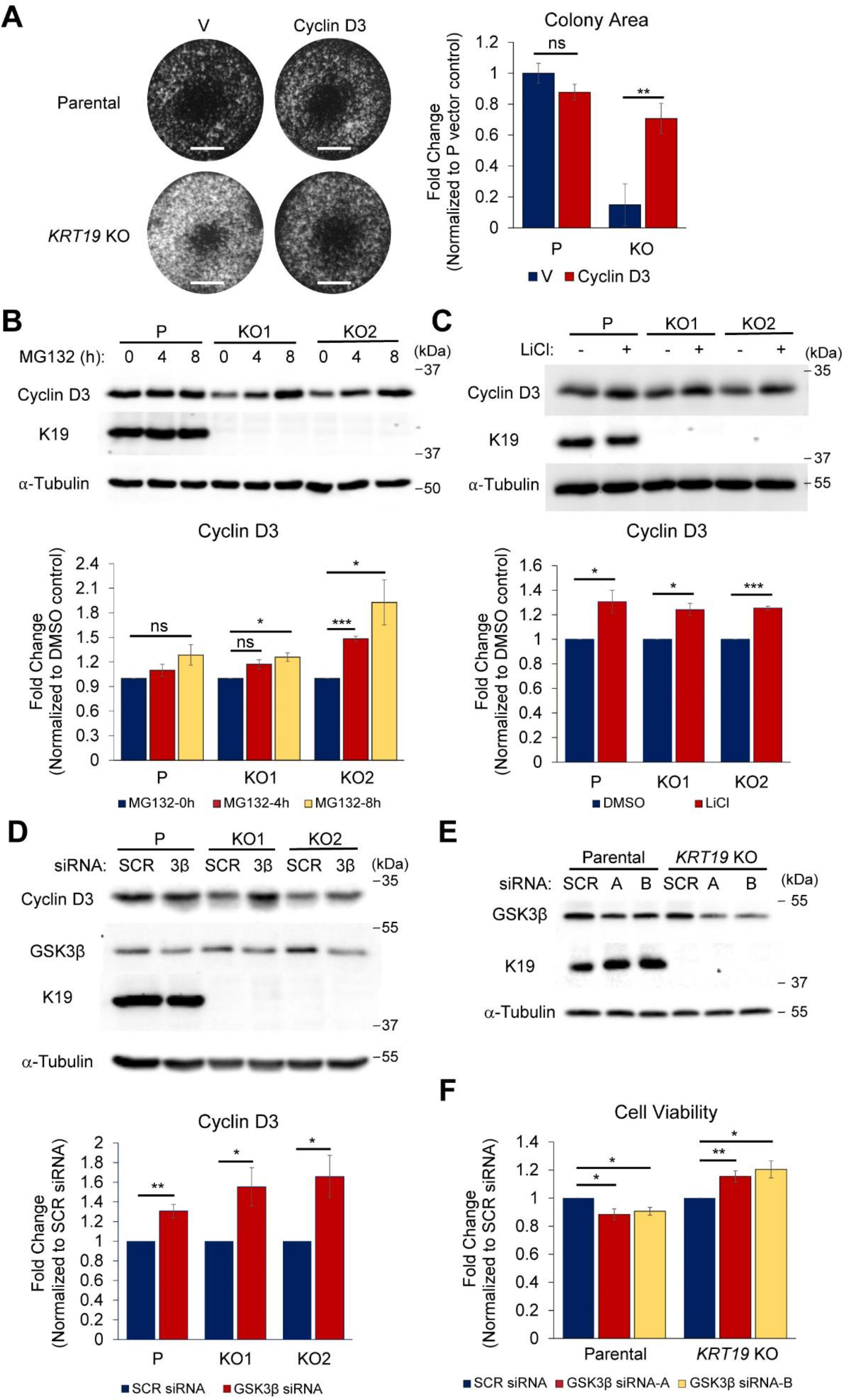
GSK3β downregulates cyclin D3 in *KRT19* KO cells. A) Colony formation assay was performed in parental (P) and *KRT19* KO (KO) cells transiently transfected with vector control or cyclin D3. Colony area normalized to parental cells transfected with vector control are shown as mean ± SEM. N = 3. Bar, 5 mm. B) Whole cell lysates of parental (P) and *KRT19* KO (KO1 and KO2) cells treated with 10 nM MG132 for the indicated time periods were harvested, and immunoblotting was performed with antibodies against the indicated proteins. Signal intensities of cyclin D3 normalized to the α-Tubulin loading control and 0 h controls are shown as mean ± SEM. N = 4. C) Whole cell lysates of parental (P) and *KRT19* KO (KO1 and KO2) cells treated with 10 mM LiCl (+) or DMSO vehicle control (−) for 8 h were harvested, and immunoblotting was performed with antibodies against the indicated proteins. Signal intensities of cyclin D3 normalized to the α-Tubulin loading control and DMSO controls are shown as mean ± SEM. N = 4. D) Whole cell lysates of parental (P) and *KRT19* KO (KO1 and KO2) cells transfected with GSK3β (3β) or scrambled (SCR) siRNA for 48 h were harvested, and immunoblotting was performed with antibodies against the indicated proteins. Signal intensities of cyclin D3 normalized to the α-Tubulin loading control and SCR siRNA transfected controls are shown as mean ± SEM. N = 6. E) Whole cell lysates of parental and *KRT19* KO cells transfected with two different GSK3β (A and B) or scrambled (SCR) siRNA for 72 h were harvested, and immunoblotting was performed with antibodies against the indicated proteins. F) MTT assays were performed on parental and *KRT19* KO cells transfected with two different GSK3β (A and B) or scrambled (SCR) siRNA for 72 h. The absorbance at 570 nm of cells with GSK3β knockdown was normalized to that of its scrambled siRNA control to calculate cell viability. Cell viability normalized to SCR siRNA transfected controls are shown as mean ± SEM. N = 5. *P < 0.05, **P < 0.01, ***P < 0.001, and ns: not significant.

Since proteasomal degradation of cyclin D3 has been reported to be induced by its phosphorylation by GSK3β (Naderi, 2004) and p38^SAPK2^ (Casanovas *et al.*, 2004), their roles in cyclin D3 degradation in *KRT19* KO MCF7 cells were tested. Inhibiting p38 using two different p38 inhibitors SB202190 or SB203850 had little to no significant effects in cyclin D3 levels when compared to vehicle controls (Figure S3). However, inhibiting GSK3β with lithium chloride (LiCl) increased cyclin D3 levels in both parental and *KRT19* KO cells (Figure 1C). This was confirmed with GSK3β siRNA, as cyclin D3 protein levels in *KRT19* KO cells were significantly increased when normalized to levels of scrambled siRNA-treated cells (Figure 1D). In addition, GSK3β knockdown using two different siRNA increased proliferation of *KRT19* KO cells, whereas slight but statistically significant reductions were observed in parental cells (Figure 1E, F). Altogether, these data suggest that GSK3β is the culprit for reduced cyclin D3 levels and compromised proliferation rates of *KRT19* KO cells.

### Activating GSK3β facilitates cyclin D3 degradation in KRT19 KO cells

To elucidate the role of GSK3β in K19-dependent stability of cyclin D3, forskolin was used. Forskolin is an adenyl cyclase activator, and forskolin-induced elevation of intracellular cAMP activates GSK3β by decreasing phosphorylation at Ser9, leading to degradation of cyclin D proteins and inhibition of cell proliferation in various settings (Musa *et al.*, 1999; Naderi *et al.*, 2000, 2004). Although 8 h treatment of 100 μM forskolin failed to alter cyclin D3 levels in parental cells, it resulted in significantly decreased levels of cyclin D3 in *KRT19* KO cells (Figure 2A). The requirement of K19 in inhibiting forskolin-induced decrease of cyclin D3 levels was confirmed when expression of GFP-tagged K19 in *KRT19* KO cells maintained cyclin D3 levels upon forskolin induction (Figure 2B). Next, GSK3β activity was examined by monitoring phospho-GSK3β levels at Ser9 (pGSK3β). Following forskolin treatment, lower pGSK3β in *KRT19* KO cells was observed compared to parental cells (Figure 2C), suggesting that K19 suppresses GSK3β activity to protect forskolin-induced decrease in cyclin D3 levels.

**Figure 2.**
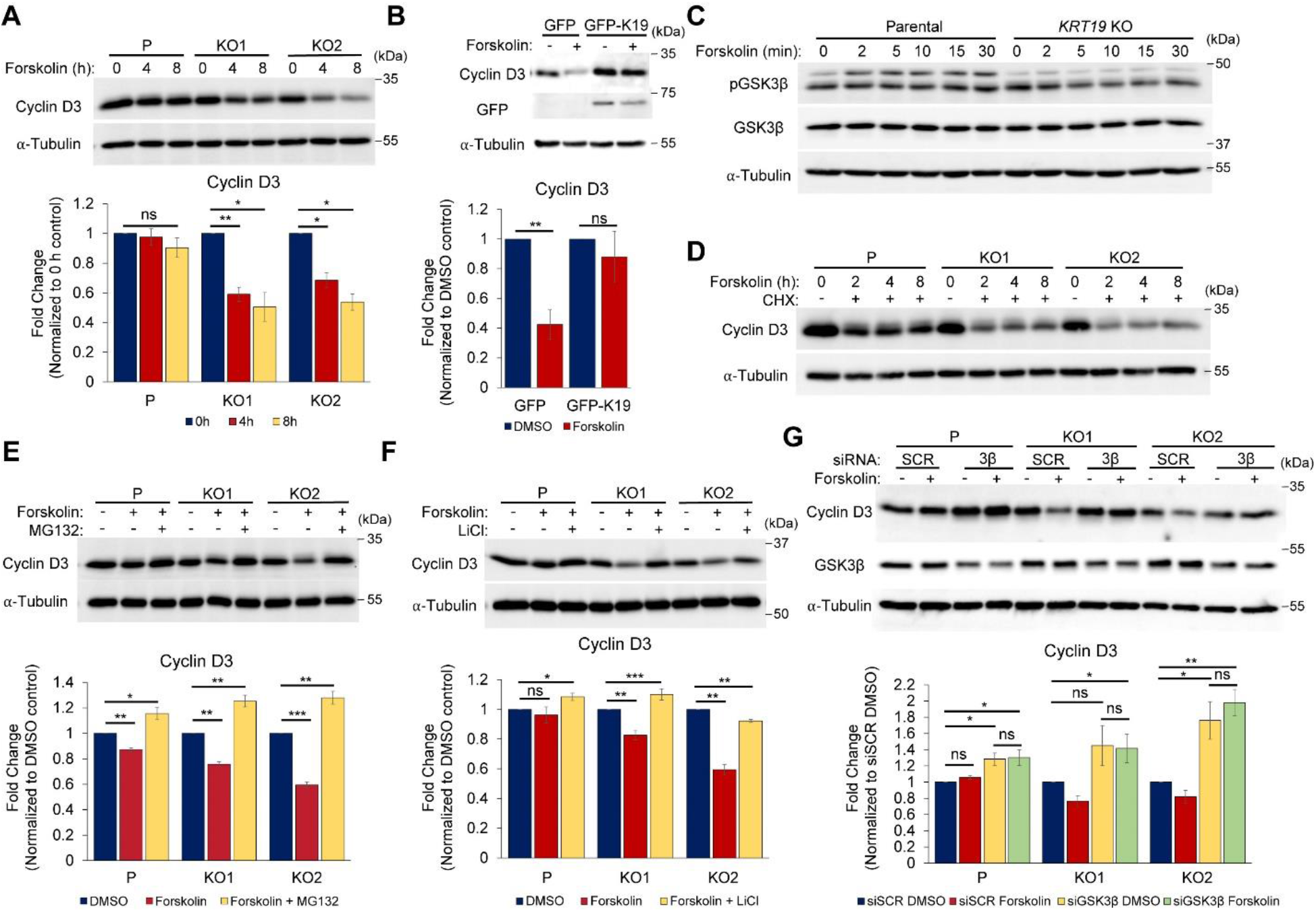
Activating GSK3β facilitates cyclin D3 degradation in *KRT19* KO cells. A) Whole cell lysates of parental (P) and *KRT19* KO (KO1 and KO2) cells treated with 100 μM forskolin for the indicated time periods were harvested, and immunoblotting was performed with antibodies against the indicated proteins. Signal intensities of cyclin D3 normalized to the α-Tubulin loading control and 0 h controls are shown as mean ± SEM. N = 4. B) *KRT19 KO* (KO2) cells stably expressing GFP or GFP-K19 were treated with 100 μM forskolin (+) or DMSO vehicle control (−) for 8 h, and whole cell lysates were harvested. Immunoblotting was performed with antibodies against the indicated proteins, and signal intensities of cyclin D3 normalized to the α-Tubulin loading control and DMSO controls are shown as mean ± SEM. N = 5. C) Whole cell lysates of parental (P) and *KRT19* KO cells treated with 100 μM forskolin for the indicated time periods were harvested, and immunoblotting was performed with antibodies against the indicated proteins. D) Parental (P) and *KRT19* KO (KO1 and KO2) cells were pretreated with 20 ng/μl of cycloheximide (CHX) or DMSO vehicle control (−) for 30 min before being treated with 100 μM forskolin for the indicated time periods. Whole cell lysates were harvested, and immunoblotting was performed with antibodies against the indicated proteins. E) Parental (P) and *KRT19* KO (KO1 and KO2) cells were pretreated with 10 nM MG132 (+) or DMSO vehicle control (−) for 30 min before being treated with 100 μM forskolin (+) or DMSO (−) for additional 8 h. Whole cell lysate were harvested, and immunoblotting was performed with antibodies against the indicated proteins. Signal intensities of cyclin D3 normalized to the α-Tubulin loading control and DMSO controls are shown as mean ± SEM. N = 4. F) Parental (P) and *KRT19* KO (KO1 and KO2) cells were pretreated with 10 mM LiCl (+) or DMSO vehicle control (−) for 30 min before being treated with 100 μM forskolin (+) or DMSO (−) for additional 8 h. Whole cell lysate were harvested, and immunoblotting was performed with antibodies against the indicated proteins. Signal intensities of cyclin D3 normalized to the α-Tubulin loading control and DMSO controls are shown as mean ± SEM. N = 4. G) Parental (P) and *KRT19* KO (KO1 and KO2) cells were transfected with GSK3β (3β) or scrambled (SCR) siRNA for 48 h before being treated with 100 μM forskolin for 8 h. Whole cell lysate were harvested, and immunoblotting was performed with antibodies against the indicated proteins. Signal intensities of cyclin D3 normalized to the α-Tubulin loading control and DMSO are shown as mean ± SEM. N = 5. *P < 0.05, **P < 0.01, ***P < 0.001, and ns: not significant.

Next, the role of K19 in forskolin-induced cyclin D3 degradation was tested by examining cyclin D3 levels following forskolin treatment in the presence of the protein synthesis inhibitor cycloheximide. With inhibition of protein synthesis, forskolin treatment decreased cyclin D3 levels in *KRT19* KO cells markedly compared to parental cells (Figure 2D). Also, when cells were pretreated with MG132, forskolin treatment failed to reduce cyclin D3 levels (Figure 2E). Forskolin-induced decrease in cyclin D3 levels was then confirmed to be mediated by GSK3β in *KRT19* KO cells, as inhibiting GSK3β activity using LiCl (Figure 2F) or decreasing GSK3β levels with siRNA (Figure 2G) prevented decrease in cyclin D3 levels upon forskolin treatment in *KRT19* KO cells (Figure 2F, G).

### K19 interacts with active GSK3β

To elucidate the mechanism underlying K19-dependent inhibition of GSK3β activity, K19-GSK3β interaction was assessed by co-immunoprecipitation (co-IP) (Figure 3A). Under normal growth conditions, immunoprecipitating K19 pulled down GSK3β and vice versa, demonstrating an interaction between two proteins. Then, forskolin and other small molecules were used to assess how GSK3β activity affects K19-GSK3β interaction. Activating GSK3β using forskolin or Akt inhibitor LY294002 resulted in GSK3β-K19 interaction, while treating cells with GSK3β inhibitor LiCl or phosphatase inhibitor okadaic acid failed to do so (Figure 3B). Consistent with this, constitutively active (S9A) but not wild type or kinase dead (K85A) GSK3β co-immunoprecipitated with K19 (Figure 3C), confirming requirement for GSK3β activity in K19-GSK3β interaction.

**Figure 3.**
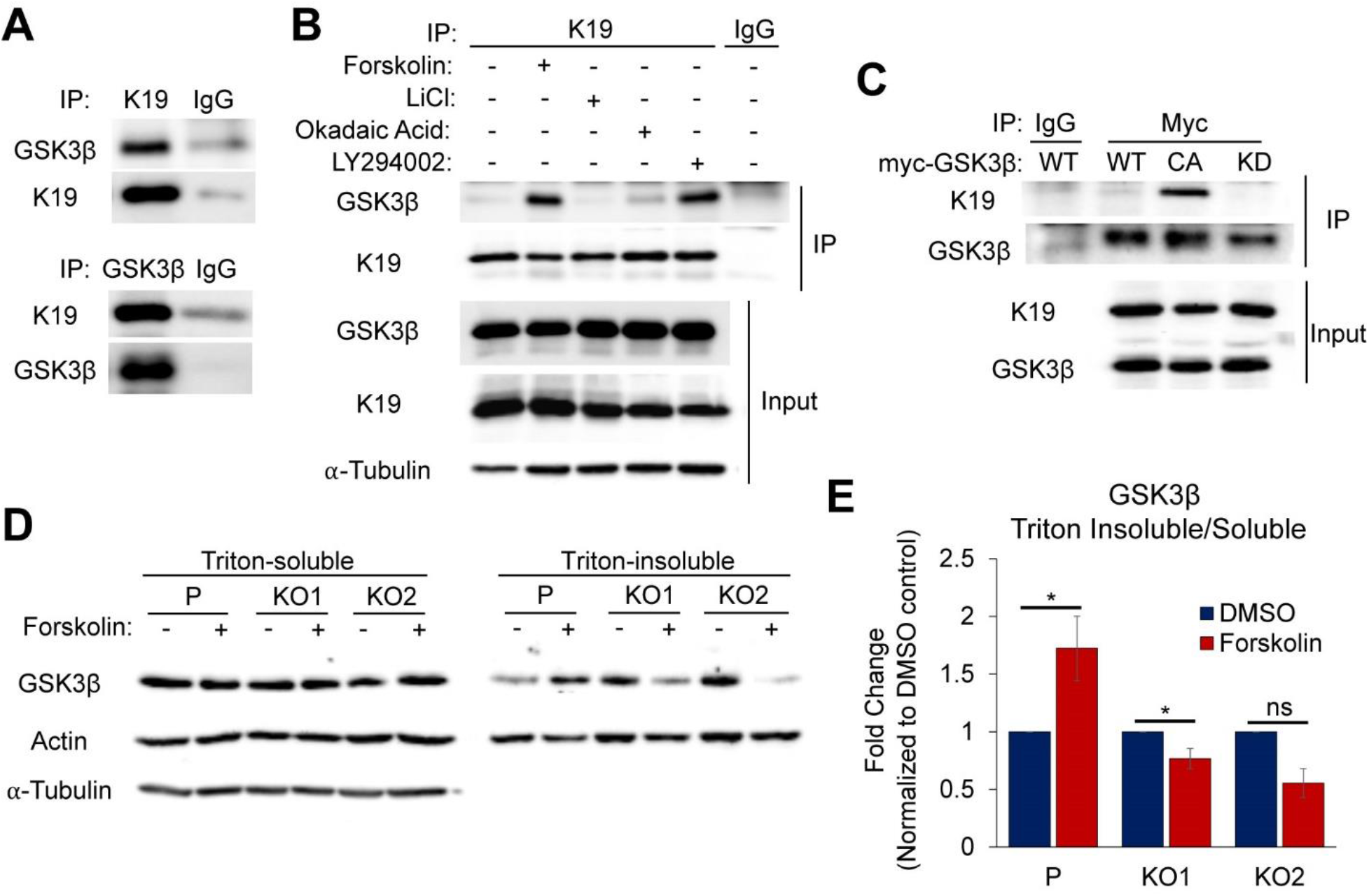
K19 interacts with GSK3β. A) Co-immunoprecipitations of K19 and GSK3β. Immunoprecipitation was performed in parental cells with anti-K19 or GSK3β antibody or IgG control. Immunoprecipitates (IP) were subjected to SDS-PAGE and immunoblotting was performed with antibodies against the indicated proteins. B) Immunoprecipitation was performed with anti-K19 antibody or IgG control in parental cells treated with 100 μM Forskolin, 10 mM LiCl, 100 nM Okadaic acid, 25 μM LY294002, or DMSO vehicle control (−) for 2 h. Immunoprecipitates (IP) and inputs were subjected to SDS-PAGE and immunoblotting was performed with antibodies against the indicated proteins. C) Immunoprecipitation was performed with anti-myc antibody or IgG control in parental cells transiently transfected with myc-tagged WT, constitutively active (CA) or kinase dead (KD) GSK3β. Immunoprecipitates (IP) or inputs were subjected to SDS-PAGE and immunoblotting was performed with antibodies against the indicated proteins. D) Parental (P) and *KRT19* KO (KO1 and KO2) cells were treated with 100 μM forskolin (+) or DMSO vehicle control (−) for 10 min. Whole cell lysates were processed for triton solubility and insolubility. Immunoblotting was performed with antibodies against the indicated proteins. E) Signal intensities of GSK3β from D) normalized to the actin loading control and DMSO controls are shown as mean ± SEM. N = 6. *P < 0.05, **P < 0.01, ***P < 0.001, and ns: not significant.

To further investigate the interaction between K19-GSK3β and the role of GSK3β activity in K19-GSK3β interaction, lysis buffers with different detergents were used to identify relative amounts of GSK3β in triton-soluble fraction *versus* keratin filament-rich triton-insoluble fraction. Following forskolin stimulation, cell lysates were first prepared using triton-based lysis buffer. Then, triton-insoluble pellets prepared using high-speed centrifugations were isolated and dissolved in urea-based lysis buffer. Forskolin stimulation did not impact GSK3β levels in triton-soluble fractions of either parental or *KRT19* KO cells but specifically increased the triton-insoluble pool of GSK3β in forskolin-treated parental cells (Figure 3D). Consistent with this, quantitation showed that there was an increased ratio of triton insoluble/soluble GSK3β only in parental cells upon forskolin stimulation (Figure 3E).

### K19 is required for the cytoplasmic localization of GSK3β

GSK3β is generally a cytoplasmic protein, but its localization is dynamic and involves continual shuttling between the nucleus and cytoplasm (Bechard and Dalton, 2009). Active GSK3β becomes accumulated in the nucleus, where GSK3β phosphorylates its nuclear targets for degradation (Bechard *et al.*, 2012). Since K19 interacts with GSK3β and inhibits GSK3β activity, the ability of K19 to regulate subcellular localization of GSK3β was tested. Immunostaining of GSK3β revealed enhanced nuclear GSK3β localization in *KRT19* KO cells upon forskolin induction, whereas forskolin-induced GSK3β localization in parental cells remained largely unchanged (Figure 4A). Biochemical subcellular fractionation of parental and *KRT19* KO cells confirmed immunostaining results. Whereas GSK3β levels in cytoplasmic fractions remained unaffected by forskolin treatment in parental and *KRT19* KO cells, a significant increase of nuclear GSK3β levels was observed specifically in *KRT19* KO cells following forskolin induction (Figure 4B, C).

**Figure 4.**
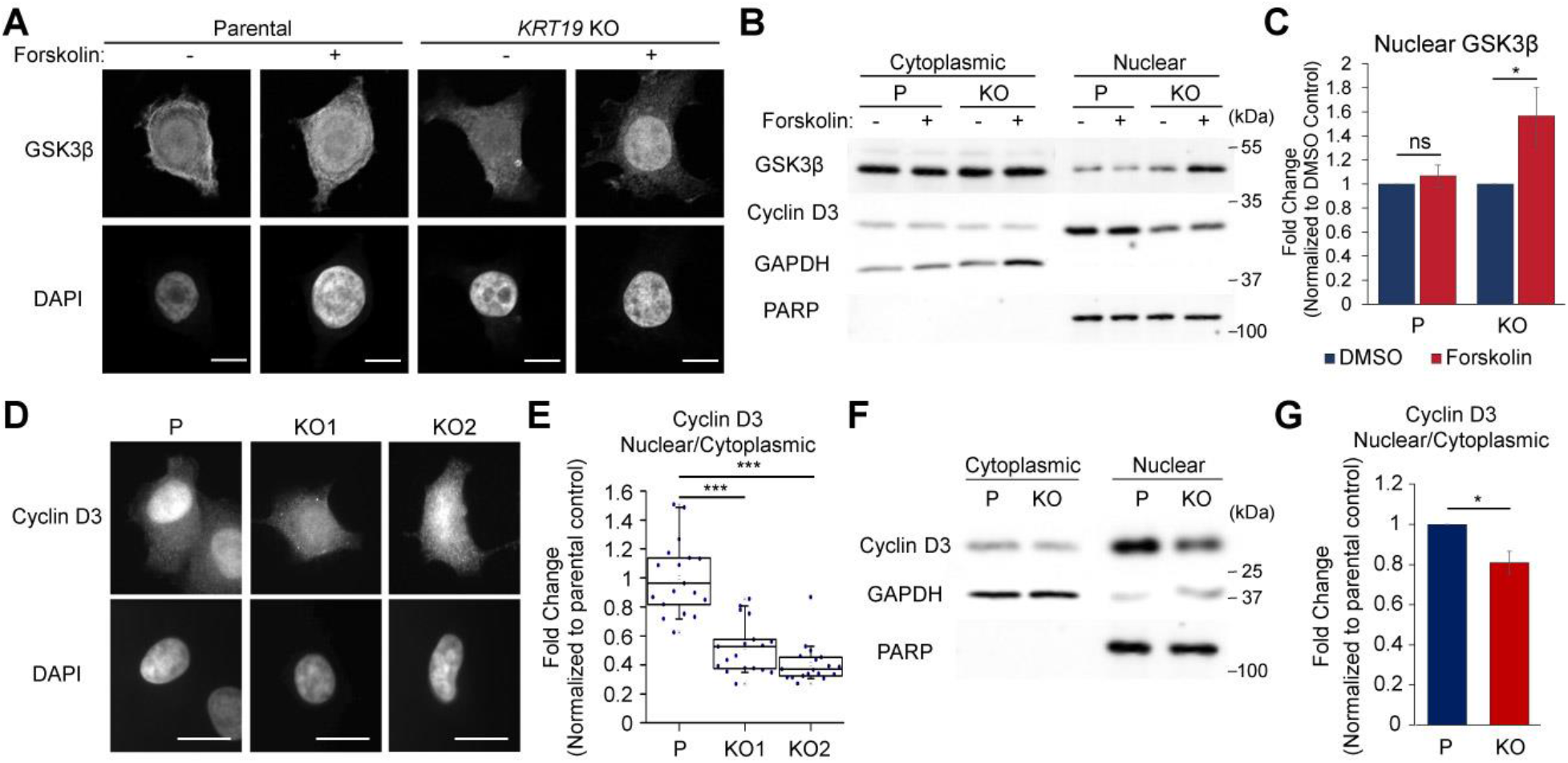
Impact of K19 on GSK3β and cyclin D3 localizations inside the cell. A) Parental and *KRT19* KO cells were treated with 100 μM forskolin (+) or DMSO vehicle control (−) for 8 h and then immunostained with anti-GSK3β antibodies. Images were obtained using a confocal microscope. Nuclei are shown with DAPI. Bar, 20 μm. B) Subcellular fractionation of parental (P) and *KRT19* KO cells treated with 100 μM forskolin (+) or DMSO vehicle control (−). Immunoblotting was performed with antibodies against the indicated proteins. PARP was used as a control for the nuclear fraction, whereas GAPDH was used for the cytoplasmic fraction. C) Signal intensities of nuclear GSK3β from B) normalized to the PARP loading control and DMSO controls are shown as mean ± SEM. N = 5. D) Parental (P) and *KRT19* KO (KO1 and KO2) cells were immunostained with anti-cyclin D3 antibody, and images were obtained using an epifluorescence microscope. Nuclei are shown with DAPI. Bar, 20 μm. E) Corrected total cell fluorescence (CTCF) of cyclin D3 from D) was quantitated and normalized to the background CTCF. Nuclear/cytoplasmic cyclin D3 levels normalized to the parental control are shown as scatter box plots with median ± maxima and minima. n = 18 cells for each cell line. F) Subcellular fractionation of parental (P) and *KRT19* KO (KO) cells. Immunoblotting was performed with antibodies against the indicated proteins. PARP was used as a control for the nuclear fraction, whereas GAPDH was used for the cytoplasmic fraction. G) Signal intensities of cyclin D3 from F) were quantitated. Nuclear/cytoplasmic ratios of cyclin D3 normalized to the parental control are shown as mean ± SEM. N = 4. * P < 0.05, **P < 0.01, ***P < 0.001, and ns: not significant.

GSK3β was reported to enhance the cytoplasmic localization of cyclin D protein (Diehl *et al.*, 1998). Immunofluorescence staining of cyclin D3 and quantitating corrected total cell fluorescence (CTCF) showed that nuclear/cytoplasmic cyclin D3 levels were significantly reduced in *KRT19* KO cells compared to those of parental cells (Figure 4D, E). Biochemical subcellular fractionation confirmed the presence of increased nuclear/cytoplasmic cyclin D3 levels in parental cells compared to *KRT19* KO cells (Figure 4F, G). These findings suggest that K19 prevents active GSK3β from shuttling into the nucleus and mediating degradation of nuclear cyclin D3.

### Identification of K19 domain required to interact with GSK3β

K19 protein contains an N-terminal head segment, very short C-terminal tail segment, and highly conserved alpha-helical central rod domain (Sharma *et al.*, 2019a). To identify the domain required for GSK3β interaction, plasmids encoding GFP-tagged wildtype (WT) or mutant with head-rod (HR), rod-tail (RT), head (H), or rod (R) domain of K19 were generated (Figure 5A) and transiently transfected into *KRT19* KO cells. GFP-tagged proteins were then pulled down using anti-GFP-conjugated beads, and the extent of co-immunoprecipitated endogenous GSK3β was measured (Figure 5B). While K19 head domain alone was not sufficient to interact with GSK3β, other domains which all included a rod domain (WT, HR, RT, and R) successfully interacted with GSK3β (Figure 5B), suggesting that the rod domain is required for the GSK3β-K19 interaction. Interestingly, while expression of WT K19 in *KRT19* KO cells rescued defects in forskolin-induced enrichment of GSK3β in the filament-rich triton-insoluble pool, RT mutant failed to do so (Figure 5C, D). Testing for the effect of GSK3β activity, forskolin-induced pGSK3β levels were increased by transfecting WT K19 compared to GFP control in *KRT19* KO cells as expected. However, RT mutant failed to increase forskolin-induced pGSK3β levels compared to GFP-expressing cells (Figure 5E, F).

**Figure 5.**
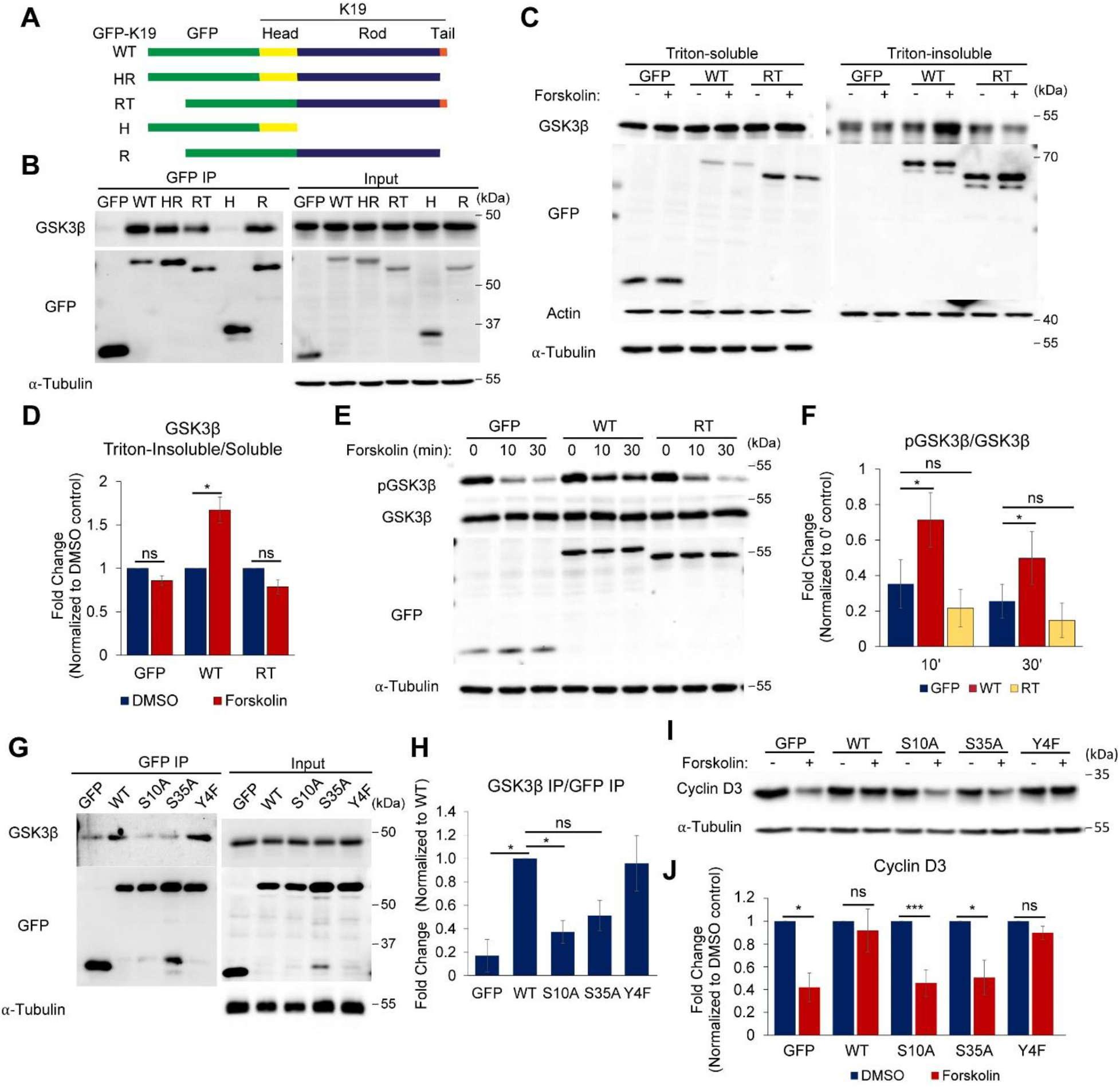
Identification of K19 domains required for GSK3β interaction. A) Schematics of K19 mutants. GFP was tagged at the N-terminus of K19 wild type (WT) and mutants. HR contains GFP fused to head and rod domains of K19; RT, GFP fused to rod and tail domains; H, GFP fused to head domain alone; and R, GFP fused to rod domain alone. B) Immunoprecipitation was performed with anti-GFP–conjugated beads in *KRT19* KO cells transiently transfected with the GFP-K19 chimeras described in A) or GFP control. Immunoprecipitates (IP) and inputs were subjected to SDS-PAGE and immunoblotting was performed with antibodies against the indicated proteins. C) *KRT19* KO cells stably expressing GFP-K19 WT (WT), GFP-K19 RT (RT) or GFP control were treated with 100 μM forskolin (+) or DMSO vehicle control (−) for 10 min. Whole cell lysates were processed for triton solubility. Immunoblotting was performed with antibodies against the indicated proteins. D) Signal intensities of GSK3β from C) normalized to the actin loading control. Triton-Insoluble/Soluble GSK3β levels normalized to DMSO controls are shown as mean ± SEM. N = 4. E) *KRT19* KO cells stably expressing GFP-K19 WT (WT), GFP-K19 RT (RT) or GFP control were treated with 100 μM forskolin for the indicated time periods. Whole cell lysates were harvested, and immunoblotting was performed with the indicated antibodies. F) Signal intensities of pGSK3β (Ser9) and GSK3β from E) were quantitated and normalized to the α-Tubulin loading control. Ratios of pGSK3β/GSK3β relative to 0’ controls are shown as mean ± SEM. N = 3. G) Immunoprecipitation was performed with anti-GFP– conjugated beads in *KRT19* KO cells transiently transfected with the indicated GFP-K19 chimeras (WT or S10A, S35A, or Y4F mutants) or GFP control. Immunoprecipitates (IP) and inputs were subjected to SDS-PAGE and immunoblotting was performed with antibodies against the indicated proteins. H) Signal intensities of GSK3β/GFP IP from G) normalized to that of WT are shown as mean ± SEM. N = 3. I) *KRT19* KO cells transiently transfected with GFP-K19 chimeras (WT or S10A, S35A, or Y4F mutants) or GFP control were treated with 100 μM forskolin for 8h. Whole cell lysates were harvested, and immunoblotting was performed with the indicated antibodies. J) Signal intensities of cyclin D3 from I) normalized to the α-Tubulin loading control and DMSO controls are shown as mean ± SEM. N = 5. *P < 0.05, **P < 0.01, ***P < 0.001, and ns: not significant.

Since the head domain of K19 was required to inhibit GSK3β activity (Figure 5E. F), we further explored to locate essential amino acids. On the head domain, there are several phosphorylation sites including Ser35 which is a major phosphorylation site on K19 (Zhou *et al.*, 1999). Keratin phosphorylation reorganizes keratin filaments and can affect their interactions with associated proteins (Eckert and Yeagle, 1990; Liao and Omary, 1996; Chung *et al.*, 2015). Therefore, we tested K19-GSK3β interaction using K19 with Y4F, S10A, or S35A mutation on the head domain. Plasmids encoding GFP-tagged WT, Y4F, S10A, or S35A K19 were transfected into *KRT19* KO cells. Immunoprecipitates of exogenous K19 were then assessed for the presence of endogenous GSK3β. Compared to WT and Y4F K19, which successfully pulled down GSK3β, GSK3β interactions with S10A as well as S35A mutants were compromised (Figure 5G, H). As K19 was required to prevent GSK3β-mediated degradation of cyclin D3, effects of K19 phospho-mutants on GSK3β-mediated degradation of cyclin D3 were also assessed. For this, cyclin D3 levels were examined following forskolin treatment in *KRT19* KO cells transfected with GFP control or GFP-K19 chimeras. While forskolin treatment did not alter cyclin D3 levels in WT or Y4F K19-expressing cells, cells expressing S10A or S35A K19 mutation showed significantly decreased levels of cyclin D3 upon forskolin treatment similar to those expressing GFP control (Figure 5I, J).

### K19 domains regulating subcellular localization of cyclin D3 and GSK3β

Since K19 head domain was required for forskolin-induced phosphorylation of GSK3β Ser9 (Figure 5E, F), the role of K19 head domain on cyclin D3 localization was examined. For this, immunostaining of cyclin D3 was done in *KRT19* KO cells stably expressing GFP control, WT, HR, or RT K19. Increased localization of nuclear cyclin D3 was observed in WT- or HR-expressing cells compared to GFP control- or RT-expressing cells (Figure 6A). Quantitating CTCF of nuclear and cytoplasmic cyclin D3 in cells revealed higher nuclear/cytoplasmic ratios of cyclin D3 in WT- or HR-expressing cells as compared to GFP control- or RT-expressing cells (Figure 6B). Biochemical subcellular fractionation also showed higher nuclear/cytoplasmic cyclin D3 levels in cells expressing WT K19 compared to those expressing GFP control or RT mutant (Figure 6C, D). Consistent with increased nuclear cyclin D3 levels, forskolin-induced nuclear accumulation of GSK3β in *KRT19* KO cells was absent in cells expressing WT K19 (Figure 6E, F). In contrast, nuclear GSK3β levels in RT K19-expressing cells mirrored those of GFP-expressing cells, suggesting that the head domain of K19 is required to prevent the nuclear localization of GSK3β. Lastly, immunostaining of cyclin D3 was performed in cells expressing GFP control or WT, S10A, or S35A K19, and CTCFs of cyclin D3 were quantitated to assess how mutations affect nuclear/cytoplasmic ratios of cyclin D3. Compared to GFP-expressing cells, nuclear/cytoplasmic cyclin D3 levels were significantly increased in cells expressing WT K19 and to a lesser extent for those expressing S35A mutant, but not for cells with S10A mutant (Figure 6G, H).

**Figure 6.**
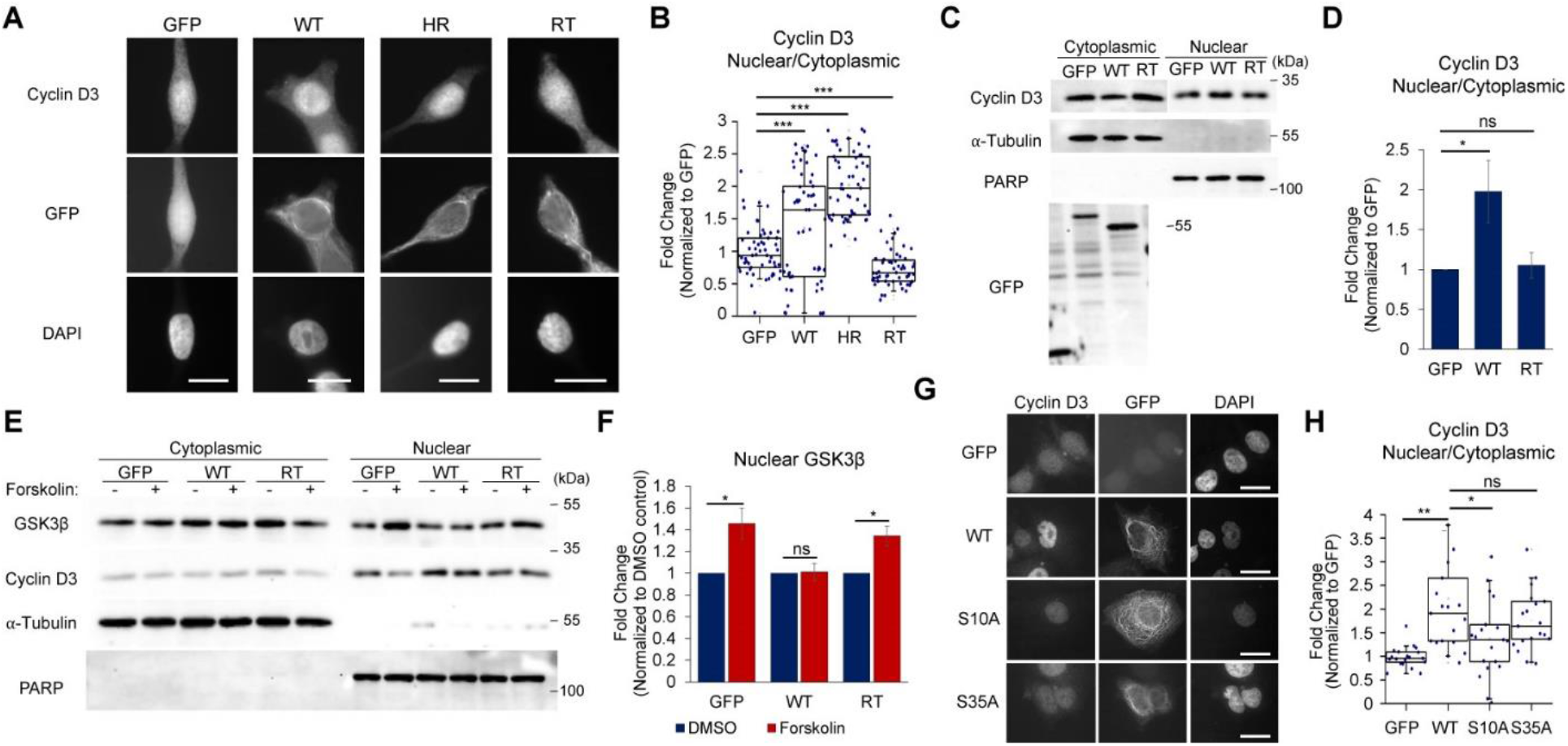
Head domain of K19 is required to maintain proper cyclin D3 and GSK3β localization. A) *KRT19* KO cells stably expressing GFP-K19 WT (WT), GFP-K19 HR (HR), GFP-K19 (RT) or GFP control were immunostained with anti-cyclin D3 and anti-GFP antibodies. Images were obtained using an epifluorescence microscope. Nuclei are shown with DAPI. Bar, 20 μm. B) CTCF of cyclin D3 from A) was quantitated and normalized to the background CTCF. Nuclear/cytoplasmic cyclin D3 levels normalized to the GFP control are shown as scatter box plots with median ± maxima and minima. n = 31 cells for each cell line. C) Subcellular fractionation of *KRT19* KO cells stably expressing GFP-K19 WT (WT), GFP-K19 RT (RT) or GFP control. Immunoblotting was performed with antibodies against the indicated proteins. PARP was used as a control for the nuclear fraction, whereas α-Tubulin was used for the cytoplasmic fraction. D) Signal intensities of cyclin D3 from C) were quantitated. Nuclear/cytoplasmic cyclin D3 levels normalized to the GFP controls are shown as mean ± SEM. N = 8. E) Subcellular fractionation of *KRT19* KO cells stably expressing GFP-K19 WT (WT), GFP-K19 RT (RT) or GFP control treated with 100μM forskolin (+) or DMSO vehicle control (−). Immunoblotting was performed with antibodies against the indicated proteins. PARP was used as a control for the nuclear fraction, whereas α-Tubulin was used for the cytoplasmic fraction. F) Signal intensities of GSK3β from E) normalized to the PARP loading control and DMSO controls are shown as mean ± SEM. N = 6. G) *KRT19* KO cells transiently transfected with GFP-K19 WT (WT), GFP-K19 S10A (S10A), GFP-K19 S35A (S35A) or GFP control were immunostained with anti-cyclin D3 and anti-GFP antibodies. Images were obtained using an epifluorescence microscope. Nuclei are shown with DAPI. Bar, 20 μm. H) CTCF of cyclin D3 from G) was quantitated and normalized to the background CTCF. Nuclear/cytoplasmic cyclin D3 levels normalized to the GFP control are shown as scatter box plots with median ± maxima and minima. n = 19 cells for each condition. *P < 0.05, **P < 0.01, ***P < 0.001, and ns: not significant.

### Impact of K19-GSK3β interaction on cell proliferation

Next, we asked whether inhibition of GSK3β activity by K19 contributes to cell proliferation. To this end, *KRT19* KO cells stably expressing various K19 mutants were tested for cell proliferation. Overexpression of WT or HR K19 increased proliferation of *KRT19* KO cells as assessed by counting cell numbers (Figure 7A), performing MTT assays (Figure 7B), and measuring confluency levels (Figure 7C) every 24 h following cell passaging. However, RT mutant lacking the head domain that was required to inhibit GSK3β activity failed to induce increased cell proliferation. This result was confirmed by performing colony formation assays (Figure 7D, E). Overexpression of WT or HR K19 showed increased colony area compared to GFP control, but RT mutant resulted in a significant decrease of colony area. To examine the effect of K19-GSK3β interaction more specifically, effects of mutations on Ser10 and Ser35 of K19 on cell proliferation were tested. While overexpression of WT K19 in *KRT19* KO cells resulted in increased colony area similar to Figure 7D, overexpressing S10A or S35A K19 was unable to induce increase in colony area (Figure 7F, G). In fact, cells overexpressing S10A K19 showed a significantly reduced area of colony.

**Figure 07.**
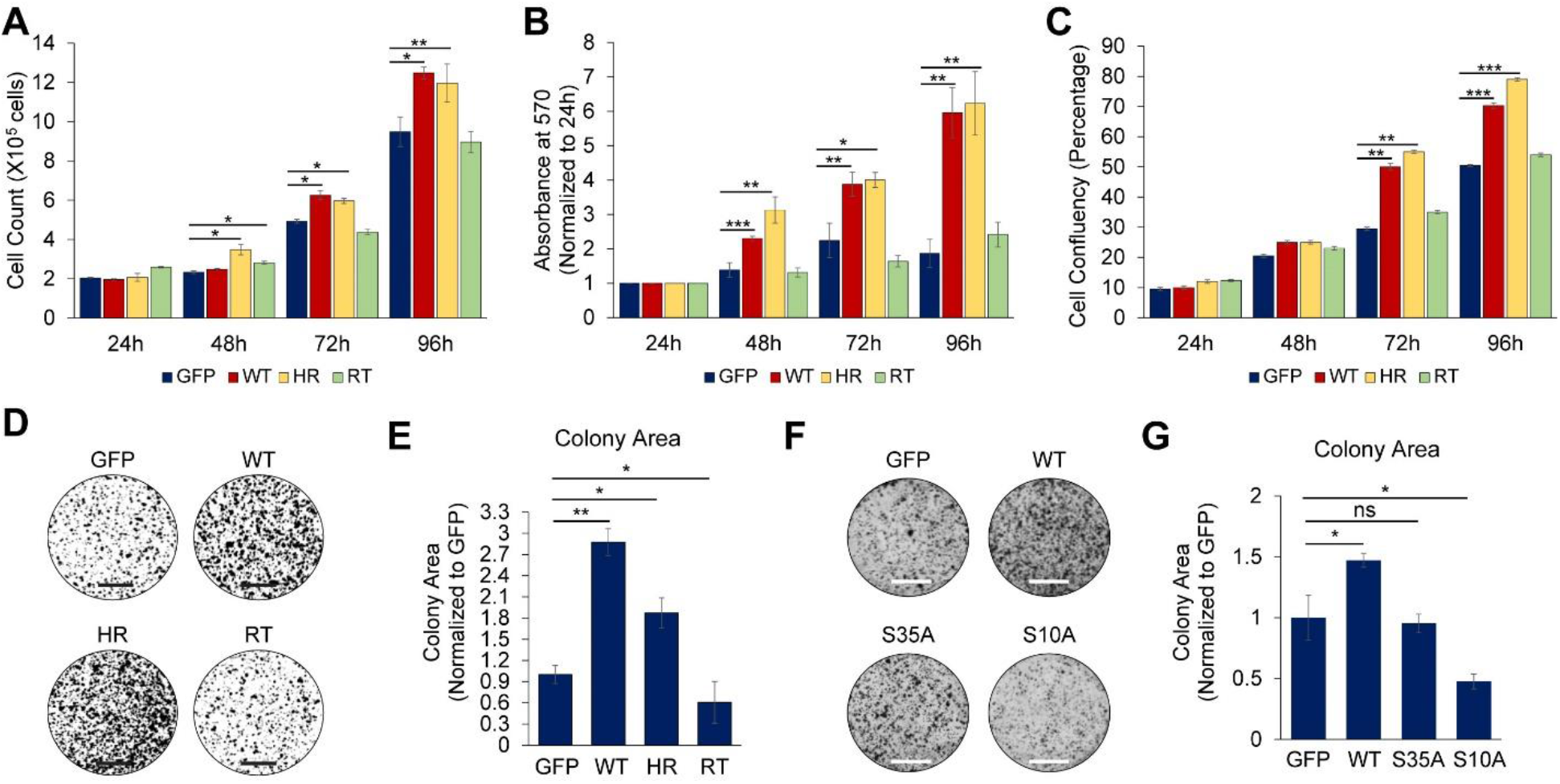
Impact of K19-GSK3β interaction on cell proliferation. Proliferation of *KRT19* KO cells stably expressing GFP-K19 WT (WT), GFP-K19 HR (HR), GFP-K19 (RT) or GFP control were assessed by A) counting cells, B) performing MTT assay and measuring the absorbance at 570 nm, and C) assessing cell confluency each day following cell plating. For B) data was normalized to absorbance at 570 nm from 24 h. For all, data from at least three biologically independent repeats are shown as mean ± SEM. D) Colony formation assay was performed in *KRT19* KO cells stably expressing GFP-K19 WT (WT), GFP-K19 HR (HR), GFP-K19 (RT) or GFP control. Bar, 5 mm. E) Colony area from D) normalized to GFP control are shown as mean ± SEM. N = 8. F) Colony formation assay was performed in *KRT19* KO *KRT19* KO cells stably expressing GFP-K19 WT (WT), GFP-K19 S10A (S10A), GFP-K19 S35A (S35A) or GFP control. Bar, 5 mm. G) Colony area from F) normalized to GFP control are shown as mean ± SEM. N = 6. * P < 0.05, **P < 0.01, ***P < 0.001, and ns: not significant.

### Inhibition of GSK3β increased sensitivity of KRT19 KO cells to CDK4/6 inhibitors

We previously found that *KRT19* KO cells showed increased resistance to CDK4/6 inhibitors Ribociclib and Palbociclib (Sharma *et al.*, 2019a). Since K19 inhibited GSK3β to stabilize cyclin D3, we examined the role of GSK3β in mediating resistance to Ribociclib and Palbociclib in *KRT19* KO cells. Cells were cultured for three days in the presence of each drug alone or in combination with GSK3β inhibitor CHIR 99021, and colony formation assay was performed to assess cell viability. The colony area of cells treated with drugs were normalized against those from cells treated with a vehicle control to calculate cell viability. As previously identified, *KRT19* KO cells were more resistant towards Ribociclib and Palbociclib, compared to the parental control (Figure 8A, B). However, co-treatment with CHIR 99021 increased the sensitivity of *KRT19* KO cells towards CDK4/6 inhibitors although co-treatment of CHIR 99021 with CDK4/6 inhibitors had no significant effects on parental cells compared to CDK4/6 inhibitors alone. Re-expressing GFP-tagged K19 in *KRT19* KO cells re-sensitized *KRT19* KO cells to CDK4/6 inhibitors (Figure 8C, D) but abrogated the effect of CHIR 99021 on how cells respond to CDK4/6 inhibitors, mimicking phenotypes of parental MCF7 cells.

**Figure 08.**
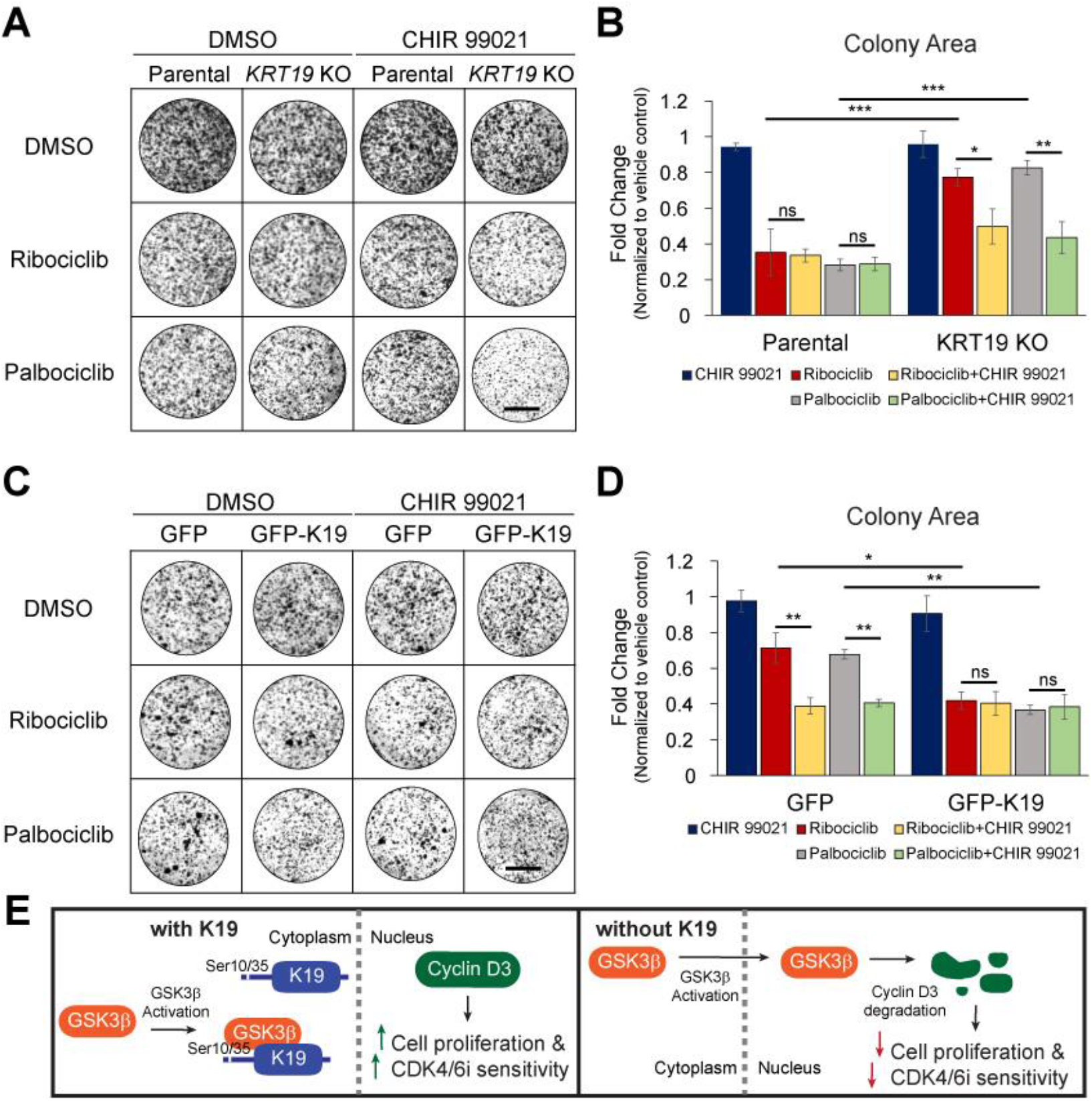
GSK3β inhibition sensitizes cells lacking K19 to CDK4/6 inhibitors. A) Colony formation assay was performed in parental and *KRT19* KO cells cultured for three days in the presence of Ribociclib or Palbociclib alone or in combination with or without CHIR 99021. Bar = 5 mm. B) Colony area from A) normalized to the parental control are shown as mean ± SEM. N = 6. C) Colony formation assay was performed using *KRT19* KO cells stably expressing GFP as control or GFP-K19 cultured for three days in the presence of ribociclib or Palbociclib alone or in combination with or without CHIR 99021. Bar = 5 mm D) Colony area from C) normalized to the parental control are shown as mean ± SEM. N = 4. *P < 0.05, **P < 0.01, ***P < 0.001, and ns: not significant. E) A model of how K19 inhibits GSK3β-mediated degradation of cyclin D3. K19 binds to GSK3β and prevents its accumulation in the nucleus. K19-GSK3β interaction requires the rod domain of K19 and activation of GSK3β. Ser10 and Ser35 of K19 also regulate K19-GSK3β interaction. Lack of K19 or introducing a GSK3β-binding deficient K19 mutant into cells increases nuclear GSK3β levels, leading to cyclin D3 degradation, decreased cell proliferation and increased sensitivity to CDK4/6 inhibitors (CDKi).

## Discussion

Overexpression of cyclin D proteins drives cancer cell proliferation (Biliran *et al.*, 2005; Zhang *et al.*, 2011), and high cyclin D levels are detected in approximately 50% of breast cancers (Barnes and Gillett, 1998; Arnold and Papanikolaou, 2005; Chi *et al.*, 2015). However, the fact that only 15-20% of breast cancers have amplification of cyclin D genes (Bartkova *et al.*, 1994; Gillett *et al.*, 1994; Dickson *et al.*, 1995; Chi *et al.*, 2015) suggests that cyclin D protein stability plays a key role in breast cancer cases. Previous evidences showed that GSK3β phosphorylation of cyclin D3 led to cyclin D3 degradation (Naderi *et al.*, 2004) and expression of kinase inactive GSKβ increased cyclin D levels (Farago *et al.*, 2005), establishing GSK3β as a negative regulator of cyclin D3. However, how GSKβ is regulated to phosphorylate cyclin D3 remained unclear.

We had previously shown that K19 is required for proliferation of MCF7 breast cancer cells (Sharma *et al.*, 2019a). This study indicates that maintenance of cyclin D3 levels underlies promotion of cell proliferation by K19, and K19-GSK3β interaction plays a key role in the process (Figure 8E). In settings where K19-GSK3β interaction was absent, GSK3β became more active, as assessed by Ser9 phosphorylation and cyclin D3 levels. Interestingly, instead of showing a preference for inactive GSK3β, K19 interacted better with active GSK3β. Therefore, K19 is likely to interact with activated GSK3β to suppresses GSK3β activity rather than interacting with inactivate GSK3β to keep it in an inactive state. However, it is still unclear how K19-GSK3β interaction impacts GSK3β activity at the molecular level. Since K19 was required to maintain phosphorylation at Ser9 when GSK3β activity was stimulated by forskolin, physical association with K19 may expose Ser9 residue of GSK3β for phosphorylation or protect phosphorylated Ser9 from phosphatases.

Instead of inhibiting GSK3β directly, K19 may be regulating GSK3β upstream regulators such as Akt to inhibit GSK3β activity. It has been reported that K19 maintains the stability of HER2, which regulates Akt (Ju *et al.*, 2015). However, the expression of HER2 is absent in MCF7 cells used in this study (Subik *et al.*, 2010). Nevertheless, other upstream regulators of GSK3β may be regulated by K19 to inhibit GSK3β activity. Because of the increasing number of keratin-interacting proteins reported so far (Sharma *et al.*, 2019a), it is likely that K19 not only binds to GSK3β but forms larger complexes with additional proteins, allowing multiple signaling proteins to communicate with one another.

Once activated by forskolin, GSK3β levels were increased in a K19 filament-rich pool of the cell. Keratin filaments have been proposed to serve as cytoplasmic scaffolds for various proteins. In particular, keratin filaments are enriched in the perinuclear region, regulating nucleocytoplasmic shuttling of proteins including 14-3-3 (Liao and Omary, 1996), hnRNP K (Chung *et al.*, 2015), β-catenin/RAC1 (Saha *et al.*, 2016), and Egr1 (Ju *et al.*, 2013). Indeed, increase in nuclear GSK3β levels upon forskolin treatment in *KRT19* KO cells suggests that physical association with K19 filaments prevents activated GSK3β from entering the nucleus. Further studies are required to investigate the exact mechanism of how K19 maintains the cytoplasmic level of GSK3β, which may involve nucleocytoplasmic shuttling of K19 itself (Bader *et al.*, 1991; Hobbs *et al.*, 2016).

While K19 head domain was required to inhibit GSK3β activity and rescue defects in cell proliferation, the rod domain was required for K19 to interact with GSK3β and the head domain by itself was not. Since the rod domain is conserved among intermediate filament proteins and critical for filament formation and stability (Zhou *et al.*, 2010; Bray *et al.*, 2015; Jacob *et al.*, 2018), it may be that filament assembly is required for K19 to initiate interaction with GSK3β. However, when cyclin D3 levels and cell proliferation were tested, the presence of the rod domain in rod-tail chimera, S10A, or S35A mutant could not rescue defects associated with the absence of K19. Therefore, while K19-GSK3β interaction may require the rod domain, it seems that the head domain harbors the ability to regulate the interaction and subsequent downstream events.

Our data points to an interesting possibility that Ser10 and Ser35 residues on the head domain of K19 may be regulated differently. While both residues were required for K19 to bind to GSK3β and maintain total cyclin D3 levels and cell proliferation, S35A mutation had milder effects compared to S10A mutation in general. At this point, the cause for the difference is unclear. Although the kinase responsible for phosphorylating Ser10 is unknown, Ser35 has been shown to be phosphorylated by Akt (Ju *et al.*, 2015). However, Akt inhibitor LY294002 increased K19-GSK3β interaction. Therefore, it may be that K19 Ser35 regulates K19-GSK3β interaction either in a mechanism independent of its phosphorylation or through phosphorylation by a kinase other than Akt. Also, S35A mutation resulted in altered filament dynamics of K19 unlike S10A mutation but the significance of altered filament dynamics to cell proliferation is unclear (Zhou *et al.*, 1999).

There is a possibility that K19 Ser10 is phosphorylated by GSK3β. On the N-terminus of K19, MTSYSYRQSS*ATSS at Ser10 matches the consensus GSK3β substrate sequence Ser/Thr-X-X-X-pSer/Thr where pSer/Thr is called a priming site that is Ser/Thr pre-phosphorylated by another kinase 4 or 5 amino acids C-terminal to the GSK3β target site (Sutherland, 2011). Also, GSK3β interaction with its substrate typically occurs when the N-terminal domain of GSK3β recognizes the phosphorylated priming site (Beurel *et al.*, 2015). Therefore, Ser14 may serve as a priming site to induce phosphorylation of Ser10 by GSK3β and thus could be involved in K19-GSK3β interaction as well. In addition, K19 Ser10 itself is 5 amino acids C-terminal to another serine, Ser5, suggesting that Ser10 could be a priming site for Ser5. This would explain why K19 S10A mutant failed to interact with GSK3β. Related to this, GSK3β was shown to phosphorylate the head domain of another intermediate filament protein desmin in hearts (Agnetti *et al.*, 2014) and muscles (Aweida *et al.*, 2018). Also, keratin filament disassembly was regulated by a GSK3β priming kinase casein kinase-1 in colon cancer cells (Kuga *et al.*, 2013). These previous reports further support the idea that K19 may be phosphorylated by GSK3β.

Our previous study showed that decreased levels of cyclin D3 in *KRT19* KO cells correlated with resistance to CDK4/6 inhibitors (Sharma *et al.*, 2019a). In this present study, GSK3β was found to be responsible for facilitating cyclin D3 degradation in the absence of K19, and the treatment of GSK3β inhibitor increased the sensitivity of *KRT19* KO cells but not parental cells to CDK4/6 inhibitors. These results suggest the high cyclin D3 levels might be responsible for making cells more sensitive to the CDK4/6 inhibitors. Indeed, cells with “D-Cyclin Activating Features (DCAF)” including elements regulating cyclin D3 expression were found to be sensitive to CDK4/6 inhibition (Gong *et al.*, 2017). Our study shows that GSK3β inhibition can be used to sensitize tumors resistant to CDK4/6 inhibitors to improve the therapeutic efficacy of CDK4/6 inhibitors. In summary, the novel mechanism presented here offers an opportunity to improve therapeutic strategy to combat cancer based on K19 expression.

## Material and Methods

### Plasmids, siRNA

Plasmids Tag5Amyc-GSK3β WT (GSK3β with C-terminal myc tag in the CMV promoter-containing pCMV-Tag 5A vector backbone, plasmid #16260), Tag5Amyc-GSK3β S9A (constitutively active GSK3β, plasmid #16261), and Tag5Amyc-GSK3β K85A (Kinase dead GSK3β, plasmid #16262) were from Addgene (Watertown, MA). Cyclin D3 cDNA was cloned out of Rc/CMV cyclin D3 (CMV promoter; #10912 from Addgene) with PCR using the primers listed in Table 1 and cloned into pmCherry-C1 (CMV promoter; Takara Bio Inc.). pEGFP-C3 K19 WT, pEGFP-C3 K19 S10A, and pEGFP-C3 K19 S35A were generated using plasmids pMRB10-K19 WT, pMRB10-K19 S10A, and pMRB10-K19 S35A (courtesy of Dr. Bishr Omary (Zhou *et al.*, 1999)) along with the primers listed in Table 1 and the In-Fusion HD cloning system (Takara, Mountain View, CA) following the manufacturer’s instructions. For pEGFP-C3 K19 Y4F, the forward primer with a mutation shown in Table 1 was used to generate *KRT19* Y4F cDNA from pMRB10-K19 WT. pEGFP-C3 K19 mutants (Head-Rod, Rod-Tail, Head, and Rod) were generated from pEGFP-C3 K19 WT using the primers listed in Table 1 with the In-Fusion HD cloning system (Takara). GFP and GFP-tagged K19 WT and mutantconstructs were cloned from pEGFP-C3 K19 constructs into pLenti CMV Hygro (plasmid #17484. Addgene) using the primers listed in Table 1 with the In-Fusion HD cloning system (Takara). To silence *GSK3B* expression, siRNA oligo duplex of Locus ID 2932 (SR301979) and scrambled negative control siRNA Duplex siRNA (SR30004) were purchased from Origene (Rockville, MD).

**Table 1.**
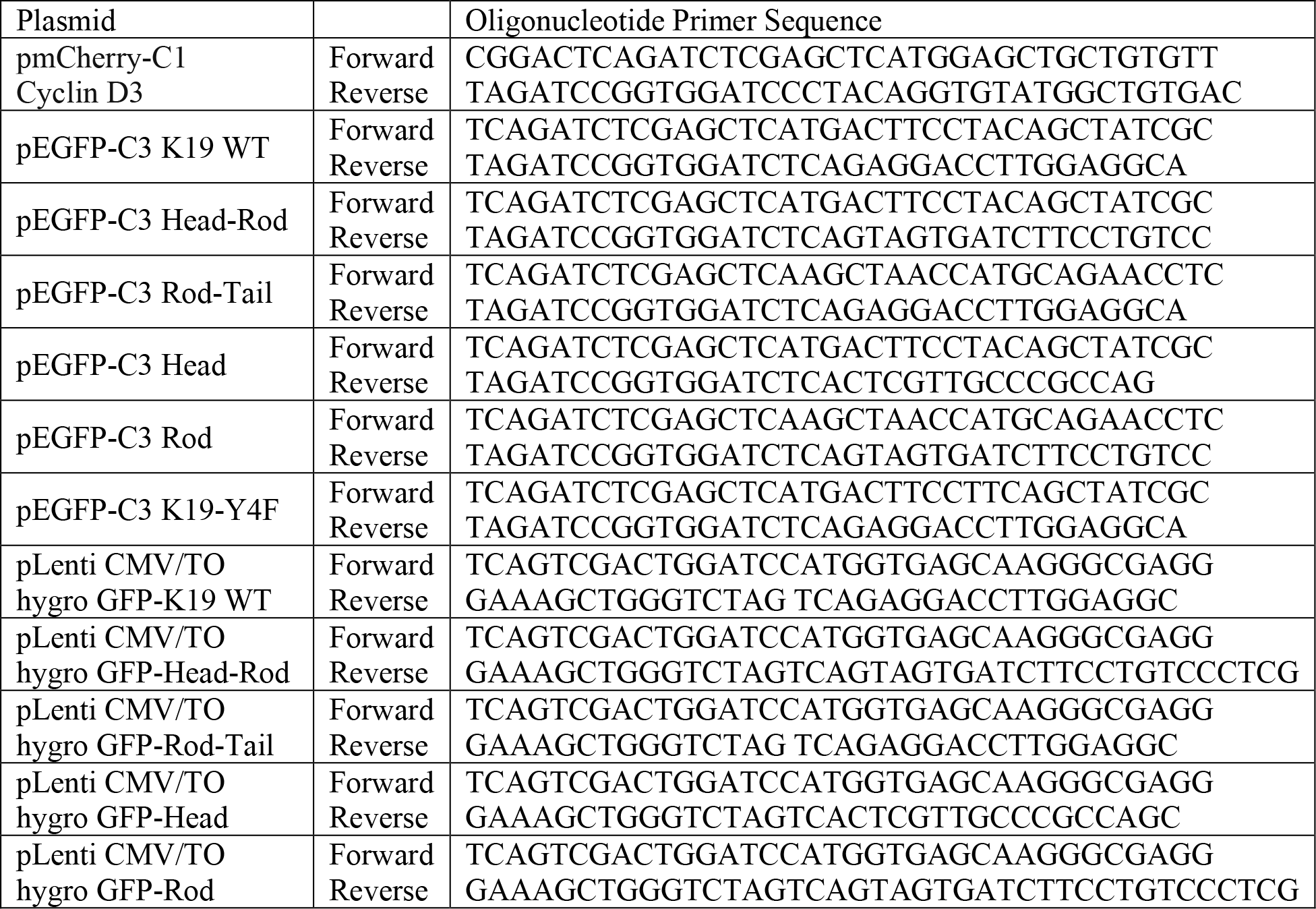
List of primers used to construct plasmids described in experimental procedures

### Antibodies and other reagents

The following antibodies: K19 (A53-B/A2), GSK3β (E-11), pGSK3β (F-2), PARP-1 (F-2), β-actin (C4), and GAPDH (0411) were from Santa Cruz Biotechnology (Santa Cruz, CA); Cyclin D3 (DCS22), and GSK3β (D5C5Z) were from Cell Signaling Technology (Danvers, MA); GFP (3H9) and RFP (5F8) were from Chromotek (Islandia, NY); Cyclin D3 (26755-1-AP) was from Proteintech (Rosemont, IL); and anti-tubulin (12G1), anti-GFP (1D2), and c-Myc (9E10) were from DSHB (Iowa City, IA). Forskolin was from ApexBio (#B1421) (Houston, TX); LiCl (# EM-LX0331) and DMSO from VWR Life sciences (Carlsbad, CA); LY294002 was from AmBeed, (#A133122) (Arlington Hts, IL); SB 203580 (#S-3400) and SB 202190 (S-1700) were from LC Laboratories (Woburn, MA); Okadaic Acid (459616) was from EMD Millipore (Billerica, MA); cycloheximide (#66-81-9) was from Sigma-Aldrich (St Louis, MO); and MG132 (S2619) was from Selleck Chemicals.

### Cell culture

MCF7 *KRT19* KO cells that were generated using the CRISPR/Cas9 system (Sharma *et al.*, 2019a) and isogenic parental control cells (ATCC, Manassas VA) were grown in Dulbecco’s Modified Essential Medium (VWR Life Science) containing 10% fetalgro bovine growth serum (RMBIO, Missoula, MT), and 100 units/ml penicillin-100 μg/ml streptomycin (GE Healthcare, Logan UT) at 37 °C in 5% CO_2_. MCF7 cells were authenticated to be an exact match (100%) of MCF7 (ATCC, HTB-22) cells using short-tandem repeat profiling service performed by ATCC (date performed: 12/28/18). For cells stably expressing pLenti CMV/TO hygro vector, GFP, or GFP-K19 chimeras, the medium was supplemented with 100 μg/ml hygromycin (#H-270-5; Goldbio, St Louis MO). For serum stimulation, cells grown in 0.1% fetalgro bovine growth serum-containing media for 48 h was stimulated with 10% fetalgro bovine growth serum-containing media or left untreated for 24 h. For forskolin treatment, cells were serum starved for 48 h and then treated with either 100 μM of forskolin or vehicle control (DMSO) for indicated time periods. For Ribociclib, Palbociclib or CHIR 99021 treatment, cells were treated for 72 hours with indicated concentrations of drugs. For pretreatment of inhibitors (LiCl, LY294002, SB 203850, SB 202190, Okadaic acid, cycloheximide, or MG132), cells were incubated with inhibitor or vehicle control for 2 h then treated with forskolin for an additional 30 min or 8 h.

### Cells stably expressing K19 mutants

Lentiviral supernatants were generated using the pLenti plasmids as described previously (Sharma *et al.*, 2019a). Lentiviral supernatants, collected 48 h after transfection, were used to infect subconfluent MCF7 *KRT19* KO cells in three sequential 4 h incubations in the presence of 4 μg/ml polybrene (Sigma-Aldrich). Transductants were selected in hygromycin (100 μg/ml), beginning 60 h after infection.

### Transfection

Overexpression plasmids were transiently transfected using Continuum™ transfection reagent (Gemini Bio-Products, West Sacramento, CA) according to the manufacturer’s protocol. siRNAs were transiently transfected using Lipofectamine RNAiMAX transfection reagent (Life Technologies) using the manufacturer’s protocol.

### Biochemical subcellular fractionation

Subcellular fractionation was performed as described previously (Chung *et al.*, 2015). Cells were washed in 1X PBS and lysed at 4°C in ice-cold cytoplasmic extract buffer (10 mM HEPES, pH 7.9; 10 mM KCl; 1.5 mM MgCl_2_; 0.1 mM EDTA; 0.5 mM dithiothreitol; 0.4% NP-40; 1 mM phenylmethylsulfonyl fluoride; 10 mM sodium pyrophosphate; 1 μg/ml each of chymostatin, leupeptin, and pepstatin; 10 μg/ml each of aprotinin and benzamidine; 2 μg/ml antipain; 1 mM sodium orthovanadate; and 50 mM sodium fluoride). Lysates were centrifuged for 10 min at 200 *g* and 4°C, and supernatants were collected as cytosolic fractions. Pellets were washed thrice with cytoplasmic extract buffers and the incubated for 1h at 4°C in a high-salt nuclear extract buffer (20 mM HEPES, pH 7.9; 420 mM NaCl; 1.5 mM MgCl_2_; 25% [vol/vol] glycerol; 0.5 mM PMSF; 0.2 mM EDTA; 0.5 mM dithiothreitol; 1 mM phenylmethylsulfonyl fluoride; 10 mM sodium pyrophosphate; 1 μg/ml each of chymostatin, leupeptin, and pepstatin; 10 μg/ml each of aprotinin and benzamidine; 2 μg/ml antipain; 1 mM sodium orthovanadate; and 50 mM sodium fluoride). Supernatants were collected as nuclear fractions after centrifugation for 10 min at 13,800 *g* and 4°C.

### MTT assay

MTT assay was performed as described previously (Sharma *et al.*, 2019a). 1000 cells were plated into each well of 96 well plate and grown in 37 °C with 5% of CO_2_ and 95% of relative humidity condition. On the day of the experiment, cells were incubated with 0.5 mg/ml of MTT [3-(4,5-dimethylthiazol-2-yl)-2,5-diphenyltetrazolium bromide] (Alfa Aesar, Haverhill, MA) containing media for 3.5 h, and formed formazan crystals were dissolved with 150 μL of isopropyl alcohol at 4 mM HCl, 0.1% NP40. The absorbance of plate was then measured at 570 nm on a SpectraMax microplate reader (Molecular Devices, San Jose, CA) and the data results were processed on a SoftMax Pro software (Molecular Devices). Each experiment was performed at least in triplicate.

### Colony formation assay

Colony formation assay was performed as described previously (Franken *et al.*, 2006). 10,000 cells per well were plated in a 6-well plate. Cells were then grown for 72 h and at the end of the experiments, cells were fixed with 3.7% formaldehyde and stained using 0.1% crystal violet for 30 min. Colony area were calculated using ImageJ (NIH). To determine the effect of Ribociclib, Palbociclib and CHIR 99021 on cell viability, cells were treated with DMSO vehicle control, 1 μM Ribociclib, 700 nM Palbociclib, with or without 5 nM CHIR 99021 24 h after plating. Cells were then grown for 96 h, and colony formation assay was performed. Colony area from drug-treated cells were normalized to those from DMSO-treated cells to calculate cell viability.

### Cell counting for proliferation

Cell counting was performed as described previously (Sharma *et al.*, 2019a). To measure cell proliferation, 50,000 cells were initially plated on each well of six-well plates. Trypsinized cells were counted using hemacytometer after every 24 h following cell passaging.

### Preparation of cell lysates, protein gel electrophoresis, and immunoblotting

Cell lysates were prepared as described previously (Chung *et al.*, 2012; Sharma *et al.*, 2019a). Cells grown on tissue culture plates were washed with 1X PBS and lysed in cold triton lysis buffer solution (1% Triton X-100, 40 mM HEPES (pH 7.5), 120 mM sodium chloride, 1 mM EDTA, 1 mM phenyl methylsulfonyl fluoride, 10 mM sodium pyrophosphate, 1 μg/ml each of cymostatin, leupeptin and pepstatin, 10 μg/ml each of aprotinin and benzamidine, 2 μg/ml antipain, 1 mM sodium orthovanadate, 50 mM sodium fluoride). To isolate the filament-rich pool of proteins, triton-soluble vs insoluble proteins were analyzed as described previously (Blikstad and Lazarides, 1983; Chou *et al.*, 1993; Chung *et al.*, 2012; Wang *et al.*, 2016). The insoluble material following triton lysis buffer incubation was pelleted, washed thrice with triton lysis buffer, and dissolved in urea lysis buffer (6.5 M urea; 50 mM Tris (pH 7.5); 1 mM ethylene glycol tetraacetic acid; 2 mM dithiothreitol; 1 mM phenylmethylsulfonyl fluoride; 1 μg/ml each of cymostatin, leupeptin, and pepstatin; 10 μg/ml each of aprotinin and benzamidine; 2 μg/ml antipain; 50 mM sodium fluoride). For immunoblotting, cell lysates were centrifuged to remove cell debris. Protein concentration was determined using the Bio-Rad Protein Assay (Bio-Rad) with BSA as standard then were prepared in Laemmli SDS-PAGE sample buffer. Aliquots of protein lysate were resolved by SDS-PAGE, transferred to nitrocellulose membranes (0.45 μm) (BioRad, Hercules, CA) and immunoblotted with the indicated antibodies, followed by horseradish peroxidase-conjugated goat anti-mouse or goat anti-rabbit IgG (Sigma-Aldrich) and Amersham ECL Select Western Blotting Detection Reagent or Pierce ECL Western Blotting Substrate (Thermo Scientific, Hudson, NH). Signals were detected using ChemiDoc Touch Imager (Bio-Rad) or CL1500 Imaging System (Thermo Fisher Scientific). For Western blot signal quantitation, the Image Lab software (Bio-Rad) or iBright Analysis Software (Thermo Fisher Scientific) was used.

### Co-immunoprecipitation

Co-immunoprecipitation was performed as described previously (Sharma *et al.*, 2019a). Cells were washed with 1X PBS and cell lysates prepared in cold triton lysis buffer (1% Triton X-100; 40 mM HEPES (pH 7.5); 120 mM sodium chloride; 1 mM ethylene diamine-tetraacetic acid; 1 mM phenyl methylsulfonyl fluoride; 10 mM sodium pyrophosphate; 1 μg/ml each of cymostatin, leupeptin, and pepstatin; 10 μg/ml each of aprotinin and benzamidine; 2 μg/ml antipain; 1 mM sodium orthovanadate; 50 mM sodium fluoride) supplemented with 0.2% empigen for anti-K19 IP, anti-GSK3β IP, anti-myc IP or anti-GFP IP. Cell lysates were centrifuged to remove cell debris, and protein concentration was determined using the Bio-Rad Protein Assay with BSA as standard. Aliquots of cell lysate were then incubated with the indicated antibody or IgG control, and immune complexes were captured using either Protein G Sepharose (GE Healthcare) or GFP-Trap beads were from Chromotek (Islandia, NY).

### Immunofluorescence staining

Immunofluorescence staining was performed as described previously (Lam *et al.*, 2020). For immunostaining of cells in culture, cells grown on glass coverslips (VWR Life Science) were washed in phosphate-buffered saline (PBS), fixed in 4% paraformaldehyde in PBS, and permeabilized in 0.1% Triton X-100. Samples were blocked in 5% normal goat serum (NGS) in 1X PBS for overnight before staining with primary antibodies diluted in blocking buffer for 16 h followed by Alexa Fluor 488- or Alexa Fluor 594-conjugated goat anti-mouse or goat anti-rabbit secondary antibodies (Invitrogen, Carlsbad, CA.) for 1 h. 1 μg/ml of Hoechst 33342 were used to stain nuclei, and coverslips were mounted on microscope slides with mounting media containing 1,4-diaza-bicyclo[2.2.2]octane (Electron Microscopy Sciences, Hatfield, PA). Labeled cells were visualized by epifluorescence with an Olympus BX60 Fluorescence Microscope (OPELCO, Dulles, VA) using an UPlanFl 60X NA 1.3, phase 1, oil immersion objective (Olympus). Images were taken with an HQ2 CoolSnap digital camera (Roper Scientific, Germany) and Metamorph Imaging software (Molecular Devices, Sunny Vale, CA). ImageJ software version 1.51J (NIH, Bethesda, MD) was used to process images. For confocal images, immunostained cells were observed and their images were captured under Zeiss LSM 710 confocal microscopy (Carl Zeiss). The captured images were processed using Zen Blue software (Carl Zeiss). Corrected total cell fluorescence (CTCF) was calculated using the ImageJ software, according to previously described protocols to control for local background fluorescence and cell size (82). The following formula was used: CTCF = integrated density − (area of selected cell × mean fluorescence of background readings). Calculated cytoplasmic and nuclear CTCF scores indicate fluorescence relative to their local backgrounds.

### Graphs and statistics

All graphs in the manuscript are shown as mean ± standard error of means (SEM). For comparisons between the two data sets, Student’s t-test (tails = 2, type = 1) was used, and statistically significant p-values are indicated in figures and figure legends with asterisks (*P < 0.05, **P < 0.01, ***P < 0.001).

## Acknowledgments

We thank the Chung lab members for their support. We also thank: Dr. Pamela Tuma and Dr. Ekaterina Nestorovich of the Department of Biology for sharing instruments and statistical help, respectively; and Dr. Kevin Rulo of Department of English at The Catholic University of America for helpful comments with the manuscript.

## Author Contributions

Conceptualization, P.S. and B.M.C; methodology, P.S., V.L, S.T., S.C., J.S.S., and B.M.C.; validation, P.S. and B.M.C.; formal analysis, P.S., S.T., and B.M.C.; investigation, P.S; resources, B.M.C. and J.S.S; data curation, P.S., S.T., and B.M.C.; writing-original draft preparation, P.S. and B.M.C.; writing-review and editing, B.M.C. and P.S.; visualization, P.S. and B.M.C.; supervision, B.M.C; project administration, B.M.C.; funding acquisition, B.M.C.

## Funding and additional information

This work was supported by National Cancer Institute grant R15CA2113071 (B.M.C.) and National Eye Institute grant P30EY001765 (J.S.S).

## Supplementary information

**Figure S1.**
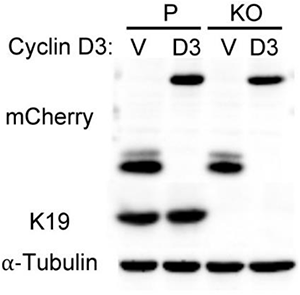
Cyclin D3 overexpression in parental and *KRT19* KO cells. Whole cell lysates of Parental (P) and *KRT19* KO (KO) cells transiently transfected with mCherry-cyclin D3 (D3) or mCherry control (V) were harvested, and immunoblotting was performed with antibodies against the indicated proteins.

**Figure S2.**
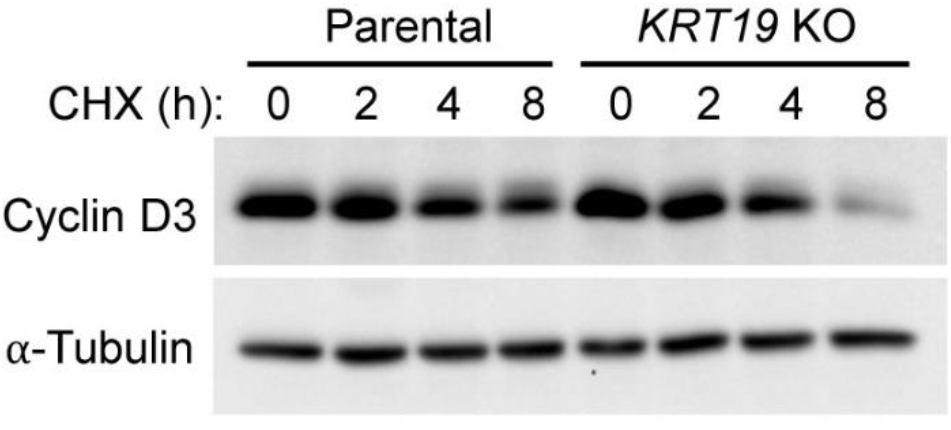
K19 regulates cyclin D3 stability. Whole cell lysates of Parental and *KRT19* KO cells treated with 20 ng/μl of cycloheximide (CHX) for the indicated time periods were harvested, and immunoblotting was performed with antibodies against the indicated proteins.

**Figure S3.**
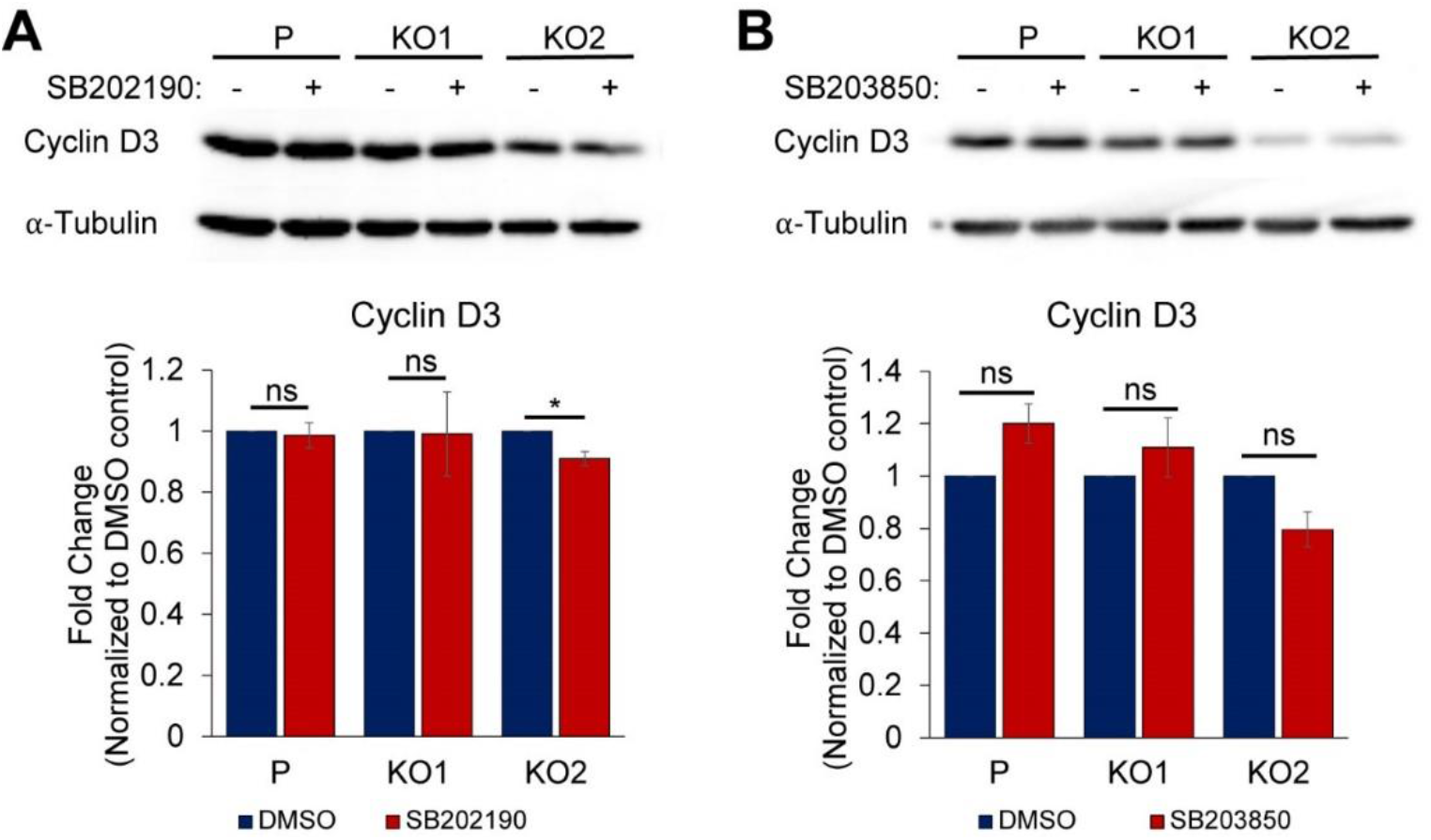
Effects of p38 inhibitors on cyclin D3 degradation in *KRT19* KO cells. Parental (P) and *KRT19* KO (KO1 and KO2) cells were pretreated with DMSO vehicle control (−) or either A) 10 μM SB202190 or B) 50 μM SB203850 for 8 h. Whole cell lysates were harvested, and immunoblotting was performed with antibodies against the indicated proteins. Signal intensities of cyclin D3 were quantified and normalized to the α-Tubulin loading control. Data normalized to DMSO controls are shown as mean ± SEM. N = 4. *P < 0.05 and ns: not significant.

## References

Agnetti, G et al. (2014). Desmin modifications associate with amyloid-like oligomers deposition in heart failure. Cardiovasc Res 102, 24–34.

Alao, JP, Stavropoulou, AV, Lam, E, and Coombes, RC (2006). Role of glycogen synthase kinase 3 beta (GSK3beta) in mediating the cytotoxic effects of the histone deacetylase inhibitor trichostatin A (TSA) in MCF-7 breast cancer cells. Mol Cancer 5, 40.

Albrecht, LV, Tejeda-Muñoz, N, Bui, MH, Cicchetto, AC, Di Biagio, D, Colozza, G, Schmid, E, Piccolo, S, Christofk, HR, and De Robertis, EM (2020). GSK3 Inhibits Macropinocytosis and Lysosomal Activity through the Wnt Destruction Complex Machinery. Cell Rep 32, 107973.

Alix-Panabieres, C, Vendrell, JP, Slijper, M, Pelle, O, Barbotte, E, Mercier, G, Jacot, W, Fabbro, M, and Pantel, K (2009). Full-length cytokeratin-19 is released by human tumor cells: a potential role in metastatic progression of breast cancer. Breast Cancer Res BCR 11, R39.

Arnold, A, and Papanikolaou, A (2005). Cyclin D1 in Breast Cancer Pathogenesis. J Clin Oncol 23, 4215–4224.

Aweida, D, Rudesky, I, Volodin, A, Shimko, E, and Cohen, S (2018). GSK3-β promotes calpain-1–mediated desmin filament depolymerization and myofibril loss in atrophy. J Cell Biol 217, 3698–3714.

Azoulay-Alfaguter, I, Elya, R, Avrahami, L, Katz, A, and Eldar-Finkelman, H (2015). Combined regulation of mTORC1 and lysosomal acidification by GSK-3 suppresses autophagy and contributes to cancer cell growth. Oncogene 34, 4613–4623.

Bader, BL, Magin, TM, Freudenmann, M, Stumpp, S, and Franke, WW (1991). Intermediate filaments formed de novo from tail-less cytokeratins in the cytoplasm and in the nucleus. J Cell Biol 115, 1293–1307.

Barnes, DM, and Gillett, CE (1998). Cyclin D1 in breast cancer. Breast Cancer Res Treat 52, 1–15.

Bartkova, J, Lukas, J, Müller, H, Lützhøt, D, Strauss, M, and Bartek, J (1994). Cyclin D1 protein expression and function in human breast cancer. Int J Cancer 57, 353–361.

Bartkova, J, Lukas, J, Strauss, M, and Bartek, J (1998). Cyclin D3: requirement for G1/S transition and high abundance in quiescent tissues suggest a dual role in proliferation and differentiation. Oncogene 17, 1027–1037.

Bechard, M, and Dalton, S (2009). Subcellular Localization of Glycogen Synthase Kinase 3β Controls Embryonic Stem Cell Self-Renewal. Mol Cell Biol 29, 2092–2104.

Bechard, M, Trost, R, Singh, AM, and Dalton, S (2012). Frat Is a Phosphatidylinositol 3-Kinase/Akt-Regulated Determinant of Glycogen Synthase Kinase 3 Subcellular Localization in Pluripotent Cells. Mol Cell Biol 32, 288–296.

Beurel, E, Grieco, SF, and Jope, RS (2015). Glycogen synthase kinase-3 (GSK3): regulation, actions, and diseases. Pharmacol Ther 0, 114–131.

Biliran, H, Wang, Y, Banerjee, S, Xu, H, Heng, H, Thakur, A, Bollig, A, Sarkar, FH, and Liao, JD (2005). Overexpression of Cyclin D1 Promotes Tumor Cell Growth and Confers Resistance to Cisplatin-Mediated Apoptosis in an Elastase-*myc* Transgene–Expressing Pancreatic Tumor Cell Line. Clin Cancer Res 11, 6075–6086.

Blikstad, I, and Lazarides, E (1983). Vimentin filaments are assembled from a soluble precursor in avian erythroid cells. J Cell Biol 96, 1803–1808.

Bray, DJ, Walsh, TR, Noro, MG, and Notman, R (2015). Complete Structure of an Epithelial Keratin Dimer: Implications for Intermediate Filament Assembly. PloS One 10, e0132706.

Casanovas, O, Jaumot, M, Paules, A-B, Agell, N, and Bachs, O (2004). P38SAPK2 phosphorylates cyclin D3 at Thr-283 and targets it for proteasomal degradation. Oncogene 23, 7537–7544.

Chi, Y, Huang, S, Liu, M, Guo, L, Shen, X, and Wu, J (2015). Cyclin D3 predicts disease-free survival in breast cancer. Cancer Cell Int 15, 89-015-0245-6. eCollection 2015.

Chou, CF, Riopel, CL, Rott, LS, and Omary, MB (1993). A significant soluble keratin fraction in “simple” epithelial cells. Lack of an apparent phosphorylation and glycosylation role in keratin solubility. J Cell Sci 105 ( Pt 2), 433–444.

Chung, BM, Arutyunov, A, Ilagan, E, Yao, N, Wills-Karp, M, and Coulombe, PA (2015). Regulation of C-X-C chemokine gene expression by keratin 17 and hnRNP K in skin tumor keratinocytes. J Cell Biol 208, 613–627.

Chung, BM, Murray, CI, Van Eyk, JE, and Coulombe, PA (2012). Identification of novel interaction between annexin A2 and keratin 17: evidence for reciprocal regulation. J Biol Chem 287, 7573–7581.

Crowe, DL, Milo, GE, and Shuler, CF (1999). Keratin 19 downregulation by oral squamous cell carcinoma lines increases invasive potential. J Dent Res 78, 1256–1263.

Dickson, C, Fantl, V, Gillett, C, Brookes, S, Bartek, J, Smith, R, Fisher, C, Barnes, D, and Peters, G (1995). Amplification of chromosome band 11q13 and a role for cyclin D1 in human breast cancer. Cancer Lett 90, 43–50.

Diehl, JA, Cheng, M, Roussel, MF, and Sherr, CJ (1998). Glycogen synthase kinase-3beta regulates cyclin D1 proteolysis and subcellular localization. Genes Dev 12, 3499–3511.

Eckert, BS, and Yeagle, PL (1990). Modulation of keratin intermediate filament distribution in vivo by induced changes in cyclic AMP-dependent phosphorylation. Cell Motil Cytoskeleton 17, 291–300.

Farago, M, Dominguez, I, Landesman-Bollag, E, Xu, X, Rosner, A, Cardiff, RD, and Seldin, DC (2005). Kinase-Inactive Glycogen Synthase Kinase 3β Promotes Wnt Signaling and Mammary Tumorigenesis. Cancer Res 65, 5792–5801.

Filipits, M, Jaeger, U, Pohl, G, Stranzl, T, Simonitsch, I, Kaider, A, Skrabs, C, and Pirker, R (2002). Cyclin D3 is a predictive and prognostic factor in diffuse large B-cell lymphoma. Clin Cancer Res Off J Am Assoc Cancer Res 8, 729–733.

Franken, NAP, Rodermond, HM, Stap, J, Haveman, J, and van Bree, C (2006). Clonogenic assay of cells in vitro. Nat Protoc 1, 2315–2319.

Giacinti, C, and Giordano, A (2006). RB and cell cycle progression. Oncogene 25, 5220–5227.

Gillett, C, Fantl, V, Smith, R, Fisher, C, Bartek, J, Dickson, C, Barnes, D, and Peters, G (1994). Amplification and overexpression of cyclin D1 in breast cancer detected by immunohistochemical staining. Cancer Res 54, 1812–1817.

Gong, X et al. (2017). Genomic Aberrations that Activate D-type Cyclins Are Associated with Enhanced Sensitivity to the CDK4 and CDK6 Inhibitor Abemaciclib. Cancer Cell 32, 761–776.e6.

Hobbs, RP, Jacob, JT, and Coulombe, PA (2016). Keratins Are Going Nuclear. Dev Cell 38, 227–233.

Jacob, JT, Coulombe, PA, Kwan, R, and Omary, MB (2018). Types I and II Keratin Intermediate Filaments. Cold Spring Harb Perspect Biol 10.

Jope, RS, and Johnson, GVW (2004). The glamour and gloom of glycogen synthase kinase-3. Trends Biochem Sci 29, 95–102.

Ju, J -h et al. (2015). Cytokeratin19 induced by HER2/ERK binds and stabilizes HER2 on cell membranes. Cell Death Differ 22, 665–676.

Ju, J-H, Yang, W, Lee, K-M, Oh, S, Nam, K, Shim, S, Shin, SY, Gye, MC, Chu, I-S, and Shin, I (2013). Regulation of cell proliferation and migration by keratin19-induced nuclear import of early growth response-1 in breast cancer cells. Clin Cancer Res Off J Am Assoc Cancer Res 19, 4335–4346.

Kabir, NN, Ronnstrand, L, and Kazi, JU (2014). Keratin 19 expression correlates with poor prognosis in breast cancer. Mol Biol Rep.

Kuga, T, Kume, H, Kawasaki, N, Sato, M, Adachi, J, Shiromizu, T, Hoshino, I, Nishimori, T, Matsubara, H, and Tomonaga, T (2013). A novel mechanism of keratin cytoskeleton organization through casein kinase I and FAM83H in colorectal cancer. J Cell Sci 126, 4721–4731.

Lam, VK, Sharma, P, Nguyen, T, Nehmetallah, G, Raub, CB, and Chung, BM (2020). Morphology, Motility, and Cytoskeletal Architecture of Breast Cancer Cells Depend on Keratin 19 and Substrate. Cytometry A, cyto.a.24011.

Liao, J, and Omary, MB (1996). 14-3-3 proteins associate with phosphorylated simple epithelial keratins during cell cycle progression and act as a solubility cofactor. J Cell Biol 133, 345–357.

Manning, BD, and Toker, A (2017). AKT/PKB Signaling: Navigating the Network. Cell 169, 381–405.

Moll, R, Franke, WW, Schiller, DL, Geiger, B, and Krepler, R (1982). The catalog of human cytokeratins: Patterns of expression in normal epithelia, tumors and cultured cells. Cell 31, 11–24.

Musa, NL, Ramakrishnan, M, Li, J, Kartha, S, Liu, P, Pestell, RG, and Hershenson, MB (1999). Forskolin Inhibits Cyclin D _1_ Expression in Cultured Airway Smooth-Muscle Cells. Am J Respir Cell Mol Biol 20, 352–358.

Musgrove, EA, Caldon, CE, Barraclough, J, Stone, A, and Sutherland, RL (2011). Cyclin D as a therapeutic target in cancer. Nat Rev Cancer 11, 558–572.

Naderi, S (2004). cAMP-induced degradation of cyclin D3 through association with GSK-3. J Cell Sci 117, 3769–3783.

Naderi, S, Gützkow, KB, Christoffersen, J, Smeland, EB, and Blomhoff, HK (2000). cAMP-mediated growth inhibition of lymphoid cells in G1: rapid down-regulation of cyclin D3 at the level of translation. Eur J Immunol 30, 1757–1768.

Naderi, S, Gutzkow, KB, Låhne, HU, Lefdal, S, Ryves, WJ, Harwood, AJ, and Blomhoff, HK (2004). cAMP-induced degradation of cyclin D3 through association with GSK-3β. J Cell Sci 117, 3769–3783.

Ohtsuka, T et al. (2016). Interaction of cytokeratin 19 head domain and HER2 in the cytoplasm leads to activation of HER2-Erk pathway. Sci Rep 6, 39557.

Petersen, OW, and Polyak, K (2010). Stem cells in the human breast. Cold Spring Harb Perspect Biol 2, a003160.

Saha, SK, Choi, HY, Kim, BW, Dayem, AA, Yang, G-M, Kim, KS, Yin, YF, and Cho, S-G (2016). KRT19 directly interacts with β-catenin/RAC1 complex to regulate NUMB-dependent NOTCH signaling pathway and breast cancer properties. Oncogene.

Sharma, P et al. (2019a). Keratin 19 regulates cell cycle pathway and sensitivity of breast cancer cells to CDK inhibitors. Sci Rep 9, 14650.

Sharma, P, Alsharif, S, Fallatah, A, and Chung, BM (2019b). Intermediate Filaments as Effectors of Cancer Development and Metastasis: A Focus on Keratins, Vimentin, and Nestin. Cells 8.

Sicinska, E et al. (2003). Requirement for cyclin D3 in lymphocyte development and T cell leukemias. Cancer Cell 4, 451–461.

Stone, MR, O’Neill, A, Lovering, RM, Strong, J, Resneck, WG, Reed, PW, Toivola, DM, Ursitti, JA, Omary, MB, and Bloch, RJ (2007). Absence of keratin 19 in mice causes skeletal myopathy with mitochondrial and sarcolemmal reorganization. J Cell Sci 120, 3999–4008.

Subik, K et al. (2010). The Expression Patterns of ER, PR, HER2, CK5/6, EGFR, Ki-67 and AR by Immunohistochemical Analysis in Breast Cancer Cell Lines. Breast Cancer Basic Clin Res 4, 35–41.

Sutherland, C (2011). What Are the *bona fide* GSK3 Substrates? Int J Alzheimers Dis 2011, 1–23.

Takano, M, Shimada, K, Fujii, T, Morita, K, Takeda, M, Nakajima, Y, Nonomura, A, Konishi, N, and Obayashi, C (2016). Keratin 19 as a key molecule in progression of human hepatocellular carcinomas through invasion and angiogenesis. BMC Cancer 16, 903.

Wang, B, Wang, Z, Han, L, Gong, S, Wang, Y, He, Z, Feng, Y, and Yang, Z (2019). Prognostic significance of cyclin D3 expression in malignancy patients: a meta-analysis. Cancer Cell Int 19, 158.

Wang, F, Zieman, A, and Coulombe, PA (2016). Skin Keratins. Methods Enzymol 568, 303–350.

Weroha, SJ, Li, SA, Tawfik, O, and Li, JJ (2006). Overexpression of cyclins D1 and D3 during estrogen-induced breast oncogenesis in female ACI rats. Carcinogenesis 27, 491–498.

Zhang, Q, Sakamoto, K, Liu, C, Triplett, AA, Lin, W, Rui, H, and Wagner, K-U (2011). Cyclin D3 compensates for the loss of Cyclin D1 during ErbB2-induced mammary tumor initiation and progression. Cancer Res 71, 7513–7524.

Zhou, Q, Snider, NT, Liao, J, Li, DH, Hong, A, Ku, NO, Cartwright, CA, and Omary, MB (2010). Characterization of in vivo keratin 19 phosphorylation on tyrosine-391. PloS One 5, e13538.

Zhou, X, Liao, J, Hu, L, Feng, L, and Omary, MB (1999). Characterization of the major physiologic phosphorylation site of human keratin 19 and its role in filament organization. J Biol Chem 274, 12861–12866.

